# Weak catch bonds make strong networks

**DOI:** 10.1101/2020.07.27.219618

**Authors:** Yuval Mulla, Mario J Avellaneda, Antoine Roland, Lucia Baldauf, Wonyeong Jung, Taeyoon Kim, Sander J Tans, Gijsje H Koenderink

## Abstract

Molecular catch bonds are ubiquitous in biology and well-studied in the context of leukocyte extravasion^1^, cellular mechanosensing^2,3^, and urinary tract infection^4^. Unlike normal (slip) bonds, catch bonds strengthen under tension. The current paradigm is that this remarkable ability enables cells to increase their adhesion in fast fluid flows^1,4^, and hence provides ‘strength-on-demand’. Recently, cytoskeletal crosslinkers have been discovered that also display catch bonding^5–8^. It has been suggested that they strengthen cells, following the strength-on-demand paradigm^9,10^. However, catch bonds tend to be weaker compared to regular (slip) bonds because they have cryptic binding sites that are often inactive^11–13^. Therefore, the role of catch bonding in the cytoskeleton remains unclear. Here we reconstitute cytoskeletal actin networks to show that catch bonds render them both stronger and more deformable than slip bonds, even though the bonds themselves are weaker. We develop a model to show that weak binding allows the catch bonds to mitigate crack initiation by moving from low- to high-tension areas in response to mechanical loading. By contrast, slip bonds remain trapped in stress-free areas. We therefore propose that the mechanism of catch bonding is typified by dissociation-on-demand rather than strength-on-demand. Dissociation-on-demand can explain how both cytolinkers^5–8,10,14,15^ and adhesins^1,2,4,12,16–20^ exploit continuous redistribution to combine mechanical strength with the adaptability required for movement and proliferation^21^. Our findings provide a new perspective on diseases where catch bonding is compromised^11,12^ such as kidney focal segmental glomerulosclerosis^22,23^, caused by the α-actinin-4 mutant studied here. Moreover, catch bonds provide a route towards creating life-like materials that combine strength with deformability^24^.

---

Here we exploit the actin-binding protein α-actinin-4 and its K225E point mutant, associated with the heritable disease kidney focal segmental glomerulosclerosis type 1^22,23^, to identify the role of catch bonds in the mechanical properties of actin networks. Actin is a key determinants of cell mechanics, together with other cytoskeletal proteins. To isolate the role of catch bonds in actin mechanics, we reconstituted actin networks from purified components. We first characterized the binding affinity of the two protein variants for actin filaments in the absence of mechanical load. Co-sedimentation of the crosslinkers with actin filaments revealed that the K255E mutant has a nearly 10-fold higher affinity (15.55 ± 0.04 μM^-1^) for actin than wild type α-actinin-4 (1.95 ± 0.04 μM^-1^, Fig. 1e and Extended Data Fig. 1). Fluorescence recovery after photobleaching measurements of crosslinker dissociation confirmed that wild type α-actinin-4 has a substantially lower bond lifetime than the mutant (Extended Data Fig. 2), consistent with prior measurements in cells^25,26^.

**Fig. 1:**
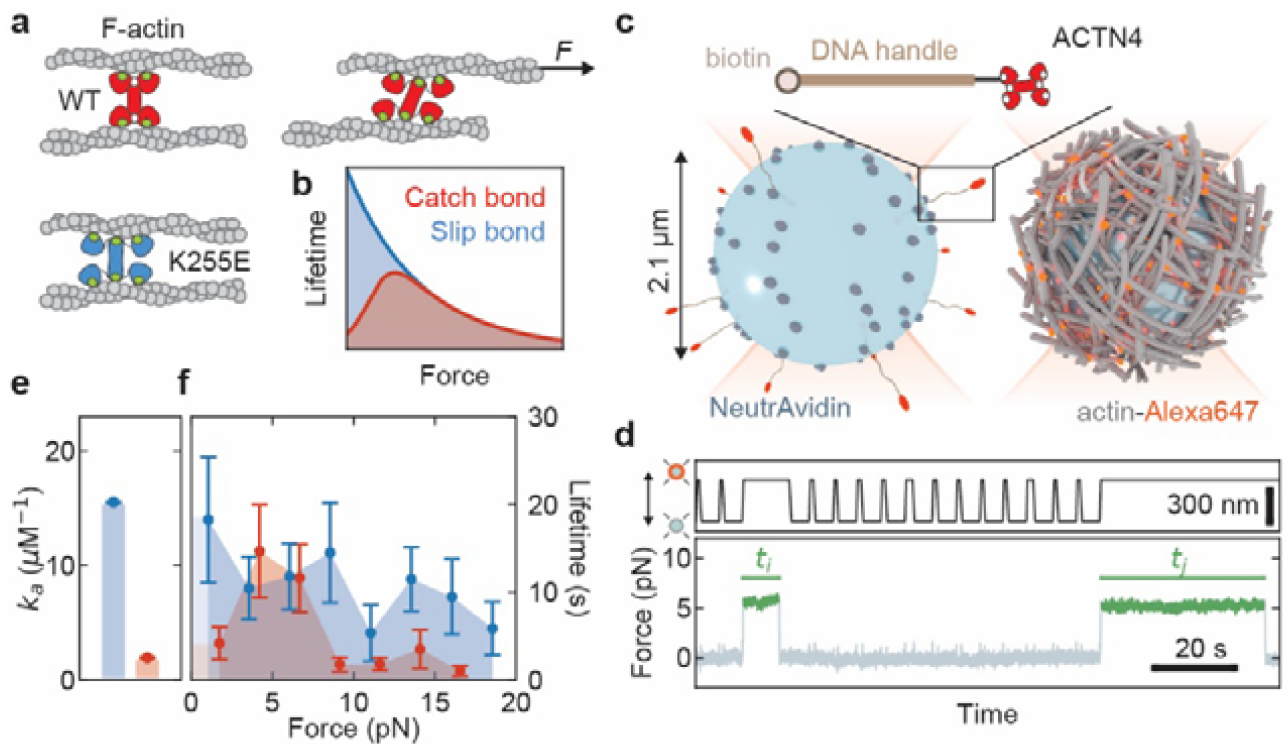
Single-molecule measurements of actin filament binding reveal catch bonding for wild type α-actinin-4 but not the K255E mutant. **a**, Each monomer of the dimeric crosslinker α-actinin-4 (red) has two weak binding sites for actin filaments (green) and one strong binding site (white) that needs to be activated by force for the wild type (WT) protein (red), whereas it is always exposed for the K255E mutant (blue). The force-induced shape transition is exaggerated for clarity. **b**, The lifetime of a catch bond first rises and then decreases with increasing force, while a constitutively active variant that acts as a regular slip bond shows a decreasing lifetime. The schematic shows a simplified limit, in which the catch and slip bond lifetimes become equal at high force. **c**, Single-molecule force spectroscopy assay, where a crosslinker-coated and an actin-coated bead are trapped using optical tweezers. **d**, Example trace illustrating the approach-and-retract protocol to establish bonds between the crosslinkers and actin filaments (top panel). An increase in the force while retracting indicates the presence of a tether (green), and the lifetime is measured until the instant the tether breaks (*t*_*i*_, *t*_*j*_, bottom panel). **e**, Actin association affinity *k*_a_ of α-actinin-4 (red) and K255E (blue) measured in a co-sedimentation assay. Error bars indicate standard error extracted from fitting the fraction of bound crosslinkers at 6 different actin concentrations assuming Michaelis-Menten kinetics (Extended Data Fig. 1a-c). **f**, Average lifetime of tethers as a function of applied force, as measured by optical tweezers (see panel d). The lifetime of wild type α-actinin-4 (red) initially rises, peaks at a force of ∼4 pN, and then decreases, as expected for a catch bond. The K255E mutant shows an overall decreasing lifetime, typical of a slip bond. Error bars indicate standard error. Affinity and force spectroscopy data were obtained at 25 °C.

## α-actinin-4 forms a weak catch bond while the K255E mutant forms a strong slip bond

Based on crystal structures, it has previously been speculated that force activates a cryptic actin-binding site of α-actinin-4, thus behaving like a catch bond^27,28^. It was furthermore proposed that the cryptic actin binding site is constitutively exposed by the K255E point mutation, increasing the binding affinity of α-actinin-4 but also abrogating its catch bond behavior (Fig. 1a, b)^13,14,25,27–30^. To directly test this idea, we tethered single α-actinin-4 molecules to polystyrene beads via DNA handles, and probed their binding to fluorescently tagged actin filaments, which fully coated another set of beads (Fig. 1c). Using optical tweezers, we trapped an α-actinin-4-coated bead and an actin-coated bead, as verified by simultaneous fluorescence imaging, (Extended Data Fig. 3b) and performed bead approach-retraction cycles. When we detected a force increase upon retraction, which indicated a binding event, we subsequently maintained the tether at a pre-set force until the force suddenly dropped to zero and the beads separated (Fig. 1d), indicating forced crosslinker unbinding. The bond lifetime for the wild type α-actinin-4 showed a load dependence consistent with catch bond behavior: short lifetimes at low loads, peaking at an intermediate load (around 4 pN), and decreasing for further increasing loads (Fig. 1f, red data). By contrast, the K255E point mutant showed slip bond behavior, with a lifetime higher than the wild type variant at low loads, consistent with the biochemical data (Fig. 1e, and Extended Data Fig. 1 and 2), and monotonically decreasing for increasing tensions (Fig. 1f, blue data). The single-molecule data provide direct proof of earlier speculations that α-actinin-4 forms weak catch bonds whilst the K255E point mutant forms strong slip bonds^13,14,25,27–30^.

## Catch bonds increase actin network strength

The observation that catch bonds are weaker than slip bonds raises the question whether they also form weaker networks. To test the strength of crosslinked actin networks, we co-polymerized actin with either crosslinker between the cone and plate of a rheometer and linearly increased the mechanical load (shear stress) in time by rotating the cone until the network ruptured, while recording the resulting network deformation (strain, Fig. 2a). We first dissected the effect of the bond lifetime on network rupturing by measuring networks crosslinked by the mutant slip bonds at either high or low temperature (25ºC for low bond lifetime and 10ºC for high bond lifetime). The high temperature was chosen such that K255E had the same bond lifetime as the wild type α-actinin-4 actin network at low temperature (Extended Data Fig. 4a-d). Consistent with intuition, we find that weaker linkers yield weaker networks (rupture stresses of 6.5 ± 0.5 Pa and 8.1 ± 1.1 Pa at 25 ºC and 10 ºC, respectively, Fig. 2b). At the same time, the weaker networks are more deformable, meaning that they reach a much larger strain before rupturing (129 ± 10% and 63 ± 4% at 25 ºC and 10 ºC, respectively, Fig. 2b). So how about the catch bonds, which have a lower bond lifetime than the mutant slip bonds at the same temperature but exhibit a different load dependence? Strikingly, networks crosslinked by the α-actinin-4 catch bonds at 10 ºC were more deformable than either of the slip bond networks (rupture strain of 221 ± 16%) yet also stronger (rupture stress of 24.5 ± 2.7 Pa, Fig. 2b).

**Fig. 2:**
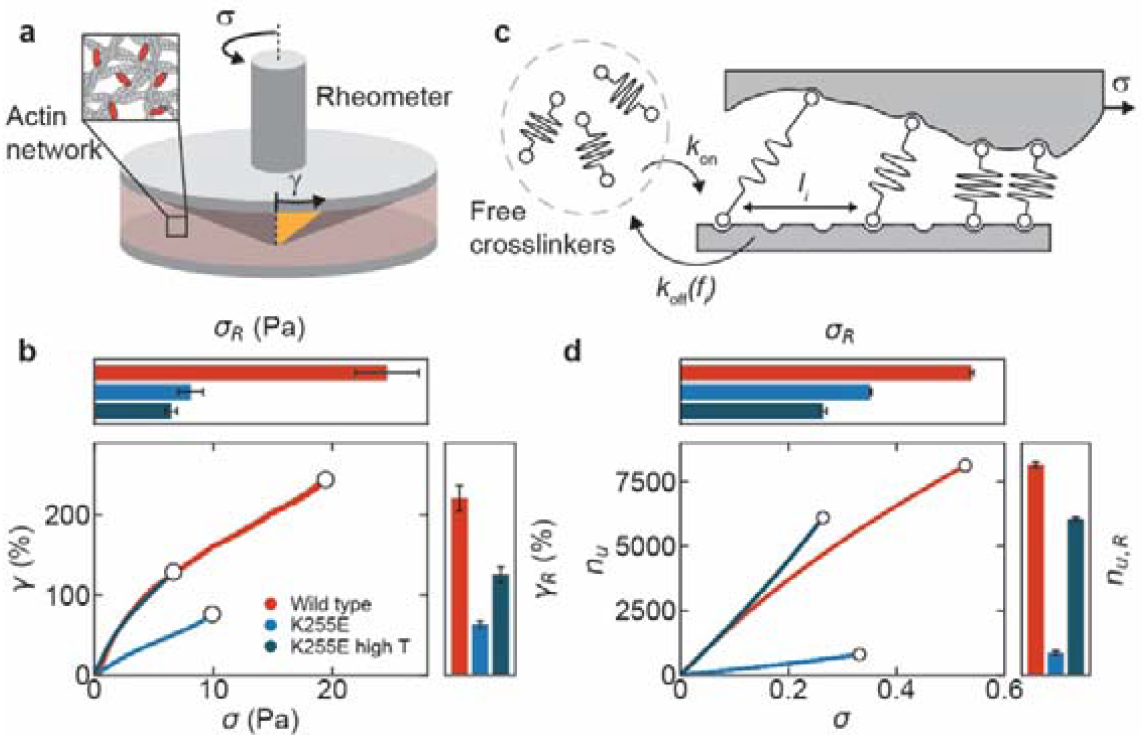
Catch bonds simultaneously enhance the mechanical strength and the deformability of cytoskeletal actin networks. **a**, Scheme of rheology experiments to characterize actin network mechanics. We measure the shear deformation *γ* of actin networks crosslinked either with α-actinin-4 or with K255E by linearly increasing the shear stress *σ* in time with a stress rate of 2.0 mPa/s. **b**, Representative examples of the shear strain *γ* as function of the shear stress *σ* for α-actinin-4 (red), K255E (blue), both at 10 °C, and for K255E at an elevated temperature (25°C, dark blue) where its lifetime matches that of wild type α-actinin-4 at 10°C (Extended Data Fig. 4c-d). The white circles indicate the rupture points (see Methods). The top panel shows the average rupture stress and the right panel the average rupture strain for each condition, with error bars representing the standard error (*N*=4 for each condition). **c**, Actin networks are modelled as 1D arrays of reversible linkers that stochastically exchange between a bound and freely diffusing state. The applied load (*σ*) linearly increases in time and is shared over all bound linkers proportionally to the distance to the nearest neighbors *l*_*i*_. **d**, The total number of unbinding events per bond *n*_*u*_ as a function of applied stress (see Methods), showing the same crosslinker dependence as the rheology experiments. The error bars in the top and right panel show the standard error (*N*=100 for each condition).

How can catch bonds escape the trade-off between strength and deformability that is inherent in normal (slip) bonds? To answer this question, we developed a minimal model where the crosslinked actin network was represented by an array of *N* reversible bonds sharing a load *σ* (Fig. 2c, see Methods and detailed explanation in the SI). We assumed nearest-neighbor load sharing (Methods Eq. 2), as it provides a simple yet accurate way to model crack initiation in viscoelastic materials, which is the rate-limiting step of rupturing (Extended Data Fig. 5)^31,32^. We used idealized Bell-Evans force-dependent unbinding kinetics to capture the catch or slip bond behavior (Fig. 1b, Methods Eq. 1)^33^, and allowed for unbound linkers to rebind at a random new location^32,34^. This bond turnover is proportional to network deformability (see SI). We chose our parameters in accordance to the force spectroscopy and biochemical data, such that the catch bonds are weaker at low force (see Extended Data Table 1 for all parameters). Strikingly, the simulations also showed that weak catch bonds collectively make networks that are stronger than slip bond networks (rupturing at nearly twice the stress, Fig. 2d), yet more deformable (with 10-fold more bond turnovers before rupturing, Fig. 2d). This difference persisted when including partially bound crosslinkers to account for the fact that α-actinin is a homodimer (Extended Data Fig. 6d, Supplementary Information). The model also confirmed the experimental observation that simply decreasing the bond lifetime while retaining a slip bond response results in weaker networks (Fig. 2d).

## Mechanism of catch bond-induced network strengthening

To identify the mechanism behind the remarkable mechanical advantage of catch bonds, we quantified the steady state distributions of the load per individual crosslinker (Fig. 3a). At a given macroscopic load, the average force per bond was only slightly higher for the catch bonds compared to the slip bonds (respectively 0.241 ± 0.003 and 0.223 ± 0.001, mean ± standard error). However, the distribution of forces for catch bonds was much narrower than for the slip bonds, meaning that slip bond networks contain substantially more bonds that bear high loads (Fig. 3a) and therefore fracture more readily. As bond load and bond-bond distance are directly proportional in our simple model (Methods Eq. 2), this means slip bond networks exhibit larger gaps than catch bond networks. To test whether suppression of large gaps by catch bonding is a potential mechanism to prevent crack initiation, we ablated adjacent bonds and simulated the network stability as a function of the gap size. Notably, gaps twice as large were required to rupture networks of catch bonds compared to slip bonds (Extended Data Fig. 5d). These findings suggest that catch bonds ‘dissociate-on-demand’ from low-stress areas thanks to their shorter lifetime at lower loads, freeing up crosslinkers that can rebind in high-stress areas and hence prevent the initiation of cracks (Fig. 3b). Simulations also showed that the mechanical advantage of catch bonds over slip bonds was indeed lost when the catch bonds are immobile (Extended Data Fig. 6c).

**Fig. 3:**
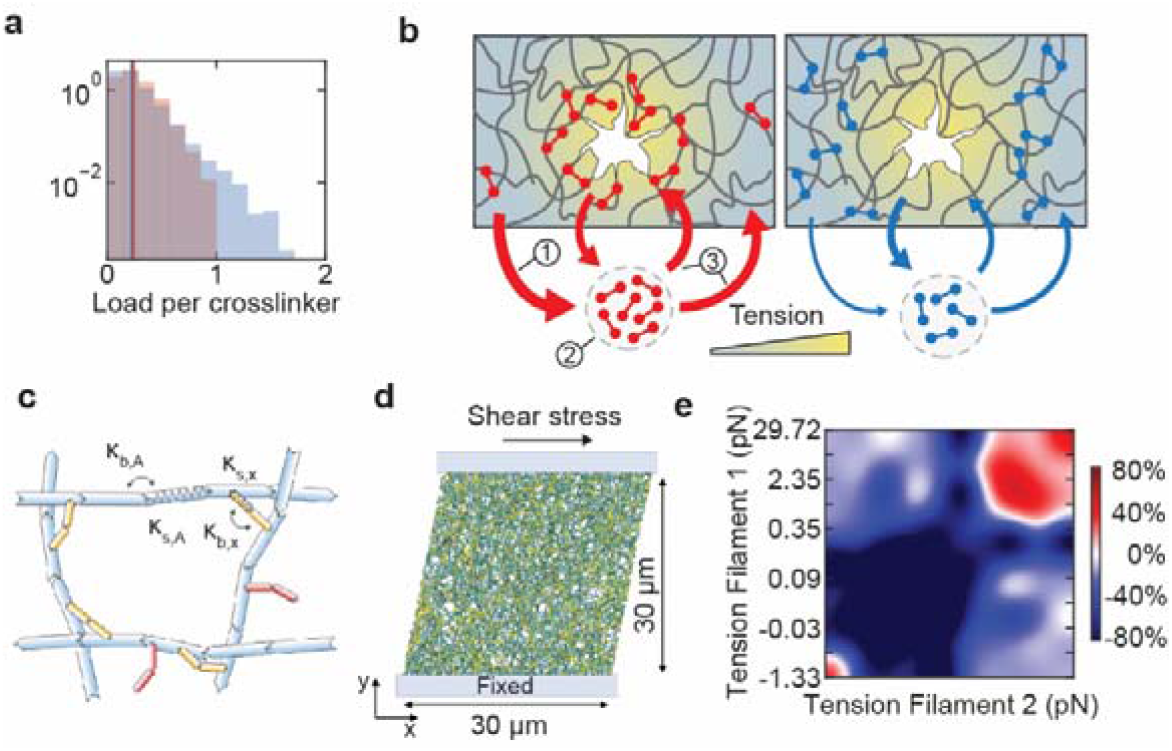
Simulations reveal that catch bonds strengthen networks by redistributing to tense areas. **a**, The distribution of forces per bond *f* measured at steady state in 1D simulations. The average bond force (vertical lines, 0.241 ± 0.003 and 0.223 ± 0.001, mean ± standard error) is larger for catch bonds (red) than for slip bonds (blue), but the force distribution of the former is much narrower: bonds carrying normalized forces higher than 1 are more than two orders of magnitude more likely for slip than for catch bonds. **b**, Self-assembly mechanism explaining the mechanical advantage of weak catch bonds (red, left) over strong slip bonds (blue, right). The thickness of the colored arrows codes for the on- and off-rate of the linkers. 1. Catch bond linkers in low tension areas rapidly unbind, increasing the pool of unbound linkers (2). As a result, there is increased binding everywhere in the network (3), at the expense of only the linkers in low tension areas. The net result is that the force distribution homogenizes, preventing crack initiation. By contrast, slip bonds preferentially localize in low-stress areas. **c**, Schematic of actin network simulations. Actin filaments (F-actins, cyan) are simplified into serially connected cylindrical segments. Crosslinkers are simplified into two arm segments connected by elastic hinges. Yellow: crosslinkers binding to two F-actins to form a functional crosslink. Red: inactive crosslinkers bound to one F-actin. Bending (κ_b_) and extensional (κ_s_) stiffnesses govern the mechanical behaviors of these segments (see Extended Data Table 2). **d**, Schematic showing how the network (30×30×1 μm) is deformed by linearly increasing shear strain by fixing the bottom of the network and displacing the top in the +x direction. **e**, Ratio of catch and slip-bond crosslinkers in color as a function of the tension in the two filaments they connect, derived from two simulations at 10 Pa stress (one with catch bonds and one with slip bonds). Color scale: percentage relative enrichment of catch bonds over slip bonds bound between a pair of actin filaments for varying tension on filament 1 (vertical axis) and on filament 2 (horizontal axis). Specifically, the color scale is defined as 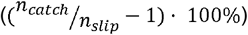 when *n*_*catch*_ > *n*_*slip*_ (shown in red) and 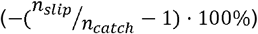 when *n*_*slip*_ > *n*_*catch*_ when *n*_*slip*_ > *n*_*catch*_ (shown in blue) where *n*_*catch*_ and *n*_*slip*_ are the number of resp. catch and slip bond linkers connecting two filaments. Compression is reported as negative tension. 10×10 tension bins are used, and the distribution was smoothed using bicubic interpolation. The tension spacing is chosen such that each bin includes 10% of the filaments (Extended Data Fig. 8b), is the same for the x- and y-axis and is only displayed on the y-axis for esthetic reasons. The simulations show that catch bonding linkers preferentially connect two high-tension filaments whereas slip bonding linkers are enriched on low-tension filament pairs.

As crosslinker rebinding is important for the mechanism, we next investigated how the mechanical advantage of catch bonding depends on the ratio between crosslinker binding and unbinding rate. We find that in case of slow binding compared to unbinding, slip bonds provide stronger networks than catch bonds (Extended Data Fig. 7a): in this regime, catch bond-induced dissociation strongly decreases the bound fraction, thereby weakening the network. By contrast, when binding is fast compared to unbinding, increased dissociation barely affects the bound fraction as crosslinkers rapidly rebind. Therefore, catch bonds provide stronger networks only when the binding rate is fast compared to the unbinding rate (Extended Data Fig. 7a), which is the relevant situation for real actin networks crosslinked by α-actinin-4 (Extended Data Fig. 1d) and also appears to be the relevant regime in cells given the strong co-localization of α-actinin-4 with the actin cytoskeleton^13,25^. To experimentally test these predictions, we performed rupturing experiments on actin networks where we tuned the unbinding rate by changing the temperature^35^. Consistent with the model’s prediction, decreasing the temperature from 25 °C to 10 °C (and hence decreasing the unbinding rate) resulted in steeper increases in the rupture stress for the α-actinin-4 catch bonds than for the K255E slip bonds (Extended Data Fig. 7b).

Our model predicts that catch bonding triggered by network stress is key to explain the increased strength of the wild α-actinin-4 crosslinkers. To test whether the loads exerted on the network were indeed sufficient to activate the catch bonds, we determined the crosslinker unbinding time from the network mechanics at different levels of shear stress, using a small oscillatory stress at different frequencies to measure the viscoelastic response time. This assay provides the characteristic network relaxation time, which is directly proportional to the crosslinker unbinding time (Supplementary Information)^36^. The network relaxation time in case of wild type α-actinin-4 crosslinkers increased with increasing shear stress, followed by a decrease at the highest stress levels, consistent with catch bonding (Extended Data Fig. 4h). For the K255E crosslinkers, the stress relaxation time was larger than for the catch bonds at low shear stress but similar at high stress (Extended Data Fig. 4h). These findings confirm the single molecule data (Fig. 1f) at the network level and show that macroscopically applied stresses above 5 Pa indeed activate strong binding for wild type α-actinin-4, whereas the K255E mutant behaves like a conventional slip bond, being strongest at small loads.

Our experiments and 1D simulations together suggest that catch bonds strengthen actin networks by being able to redistribute to tense regions under load. However, these simple simulations lack an explicit polymer network. We therefore decided to probe bond redistribution by realistic 3D-simulations of actin networks^39–42^ (Fig. 3c). These simulations contain explicit filaments and capture the main features of actin network mechanics while allowing for microscopic force measurements that are not experimentally tractable. We simulated strain ramps on networks connected by either catch or slip bonds (Fig. 3d and Extended Data Fig. 8a). It is well-known that stress is mostly carried by a small subset of tense filaments in actin networks^41^, so we analyzed how effective catch and slip bond crosslinkers were at connecting these tense filaments. Strikingly, we found that catch bonds formed up to 80% more links between pairs of highly tensed filaments than slip bonds (Fig. 3e, Extended Data Fig. 8e-f), despite binding less on average (Extended Data Fig. 8c). Furthermore, this preferential binding to stressed actin filaments increased as the bulk shear stress was raised (Supplementary Video 2). These findings directly verify the catch bond redistribution to high-stress areas as predicted by the 1D model.

## Discussion

Our work reveals a new role for catch bonds in the cytoskeleton, namely to simultaneously increase its mechanical strength and its deformability. Contrary to the common intuition that catch bonds provide strength-on-demand, our model shows that they make strong networks because dissociation-on-demand enables them to redistribute to tense areas and thus postpone crack initiation. This mechanism likely also applies to living cells, as α-actinin-4 is mobile inside the actin cortex and was recently observed to increase its bond lifetime upon mechanical stress in living cells^30^. Force sensors for α-actinin 4 analogous to those available for other proteins^43,44^ could be developed and resolved with single-molecule resolution to directly visualize the load-dependent redistribution for catch and slip bonds in both reconstituted actin networks and living cells.

Our findings also suggest a molecular mechanism to explain the low mechanical stability of kidney cells in patients afflicted by heritable disease kidney focal segmental glomerulosclerosis type 1, where α-actinin-4 carries the point mutation K255E and is known to cause podocyte fragility^22,23^. A similar mechanism may possibly apply to other diseases where loss of catch bonding leads to tissue failure, such as Von Willebrand disease 2B^11,12^. In this work we focused on the implications of catch bonds for network strength, but the generality of our model implies that this same mechanism can also apply to catch bonds in cell-matrix and cell-cell adhesions^1,2,16–20^, as dissociation-on-demand reduces friction whilst simultaneously minimizing the risk of complete cell detachment. Therefore, our results suggest that catch bonds are widespread in the cytoskeleton and at cellular interfaces to break this deformability/strength trade-off, and it would be interesting to investigate force-dependent binding of more crosslinkers and adhesins, such as filamin and IgSF CAMs.

Finally, our findings offer a cell-inspired route to create hydrogel materials that are strong yet sufficiently deformable for applications in regenerative medicine^45^. Recent years have seen a surge of theoretical and experimental work in the polymer community to create tougher hydrogels, for instance by the inclusion of stiff elements into the hydrogel^46–51^. However, these approaches have largely focused on preventing crack propagation, rather than crack initiation. Preventing crack propagation works well to absorb a finite amount of mechanical energy but offers limited advantage in case of constant stress. For future work, it would be interesting to combine mitigation strategies for both crack initiation and propagation. Synthetic analogues of catch bonds have recently been discovered and provide an excellent starting point towards highly dynamic yet strong biomimetic materials^52,53^.

## Methods

### Minimal 1D crosslinker model

To investigate the effect of molecular catch bonding on the strength of cytoskeletal filament networks, we use a computational model we recently developed to predict failure of transient networks^31,32^, using a Gillespie algorithm to model stochastic linker binding and unbinding. The detailed motivation behind the design of the model, including a discussion of its assumptions and limitations, are presented in the Supplementary Information. We consider a 1D model of *N* linkers that share an externally applied load σ (Fig. 2c). We model the effect of a force *f* on the unbinding rate *k*_off_ (inverse bond lifetime) of a bound linker *i*using the Bell-Evans equation^33^:

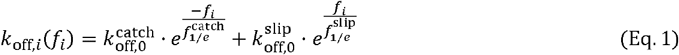

The first exponent models the catching of the weakly bound state, whereas the second exponent models the slipping of the force-activated state. We compare catch bonds with slip bonds, which do not require force-activation for strong binding 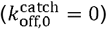, keeping all other parameters identical (see Extended Data Table 1 for the full list of parameters used for each simulation). To account for the mobility by random diffusion of the linkers after unbinding, we allow for unbound linkers to rebind at a random new location^32,34^. As the actin concentration is much larger than the crosslinker concentration both in our reconstituted networks (48 μM versus 0.48 μM, respectively) and in living cells (∼100 μM versus ∼1 μM, respectively^54^), we consider 10-fold more binding sites than crosslinkers to prevent competition for actin-binding sites. For control simulations where the linkers are immobile (Extended Data Fig. 6c), we only allow for rebinding in the same place where the crosslinker unbound^31^.

It is known that stressed networks connected by reversible bonds exhibit spontaneous crack initiation and propagation due to inhomogeneous load sharing^55^. We reproduce this rupturing behavior using a minimal model where the force per linker *f*_i_ is proportional to the global applied stress and the distance between its nearest neighbors on both sides *l*_*i*_ in 1D (Fig. 2c):

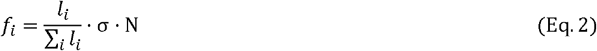

The stress is normalized by the force per crosslinker, such that 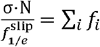. Time is normalized by the binding rate as we keep *k*_on_ =1. We use a periodic boundary condition to prevent edge effects. We initialize networks by randomly placing 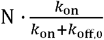 linkers (see Extended Data Table 1 for all parameters). The Supplementary Information and Extended Data Fig. 6a-b contain a more detailed discussion of the effect of network size.

### Actin network simulations

We employed an agent-based computational model based on Brownian dynamics^39–42^. In this model, the actin filaments (F-actin) and crosslinkers are simplified by cylindrical segments (Fig. 3b). The motions of the cylindrical segments are determined by the Langevin equation. Bending and extensional forces maintain angles and distances formed by the cylindrical segments near their equilibrium values, respectively. A repulsive force represents volume-exclusion effects between overlapping F-actins. The details of the model are explained in the Supplementary Information, and parameter values are listed in Extended Data Table 2. We create a crosslinked actin network in a three-dimensional thin rectangular domain (30×30×1 μm) with a periodic boundary condition only in the x-direction (Fig. 3c). The network is spontaneously assembled by dynamic events occurring on F-actins and crosslinkers; F-actin is formed by nucleation and polymerization events, and crosslinkers bind to elongating F-actin to form crosslinking points. The average filament length is ∼12 μm. It is assumed that crosslinkers unbind from F-actins at a force-dependent rate; the unbinding rate follows either the slip or catch bond behavior. After unbinding from one F-actin, crosslinkers also unbind quickly from the other F-actin, disappear, and then appear in a different location whose distance is within 5 μm from the unbinding location. During the network assembly, all crosslinkers are assumed to behave as slip bonds. After network formation, the network is deformed by applying a shear strain that linearly increases at a rate of 0.001 s^-1^. These simulations are performed with either slip or catch bond crosslinkers. The corresponding shear stress acting on the network at each strain level is measured. During shear deformation, we measure tensile forces acting on F-actins and analyze how crosslinkers are redistributed on pairs of filaments depending on tensile forces. As these simulations are strain-controlled, network yielding does not display a sudden divergence of the strain. Instead, the yielding points were determined as the first data point in which network stress decreases (dσ/dt<0). As the simulated stress/strain curves are noisy (Extended Data Fig. 8a), we convoluted the raw curves with a 60 data point time-window before determining the rupture point.

### Protein purification

Human wild type α-actinin-4 and its K255E point mutant were purified as previously described^56^. Briefly, *E. Coli* cells were transformed to express recombinant crosslinkers with a His_6_-tag. Induction was performed with 500 μM isopropyl β-D-1-thiogalactopyranoside for eight hours at 25 ^º^C. After centrifugation at 6000 g for 15 minutes, cells were resuspended in 20 mM NaCl, 5 mg/ml lysozyme and 20 mM HEPES, pH 7.8. The cells were lysed by a freeze-thaw cycle and the lysate was centrifuged at 20,000 g for 30 min. The recombinant proteins were purified from the supernatant using a QIAGEN nickel column that was first washed with 20 bed volumes of 500 mM NaCl, 25 mM imidazole, and 20 mM HEPES, pH 7.8. The recombinant proteins were eluted with 10 bed volumes of 500 mM NaCl, 500 mM imidazole, and 20 mM HEPES, pH 7.8, concentrated using Centricon filters (Millipore), and purified by gel filtration in 150 mM NaCl, 20 mM HEPES pH 7.8, and 10 mM dithiothreitol (DTT). Actin was labeled using an Alexa Fluor Labeling Kit purchased from ThermoFisher and biotin-actin was purchased from Cytoskeleton.

To ensure we compare α-actinin-4 and K255E at the same concentration in all our assays, we determined the ratio of the protein stock concentrations by measuring the intensity of the protein bands on an SDS-PAGE gel. We chose this method because, unlike UV-VIS spectrophotometry, it specifically measures the protein of interest and excludes the contribution of any contaminants. The proteins were cysteine-labeled using maleimide-activated Oregon Green at a ratio of five fluorophores for every crosslinker at room temperature for 1 h. Labeled proteins were separated from free dye molecules by gel filtration using a Superdex 200 column (GE Healthcare)^56^.

Actin was purified from rabbit psoas skeletal muscle as described in reference^34^, including the gel filtration step to remove oligomers. The concentration was determined by measuring the optical absorbance at 280 nm. Aliquots were snap-frozen and stored at -80ºC in G-buffer (2 mM tris-hydrochloride pH 8.0, 0.2 mM disodium adenosine triphosphate, 0.2 mM calcium chloride, 0.2 mM dithiothreitol) to prevent polymerization. After thawing, we stored G-actin stock samples overnight at 4ºC. The next day, we spun the sample at 120 000 g to remove any remaining aggregates. The supernatants were stored at 4 ^º^C and used within 7 days. We polymerized actin at a concentration of 48 μM (2 mg/ml) in an F-buffer consisting of 50 mM KCl, 20 mM imidazole pH 7.4, 2 mM MgCl_2_, 1 mM DTT and 0.5 mM MgATP in the presence of crosslinker at a concentration of 0.48 μM (corresponding to a molar ratio of 1/100 crosslinker/actin and on average around 1 crosslinker for every 0.5 μm length of actin filament). We verified that the networks under these conditions are isotropic and spatially uniform by confocal fluorescence imaging (Extended Data Fig. 9). Unless otherwise mentioned, all chemicals were purchased at Sigma Aldrich.

### SDS-PAGE gel protocol and quantification

SDS-PAGE gels were used to characterize and quantify purified proteins. In all cases, 20 μl sample was mixed with 20 μl InstantBlue and boiled at 95 ^º^C for 5 minutes in a closed Eppendorf vial. 30 μl of this solution was loaded onto a 4–15% Mini-PROTEAN TGX Precast Protein Gel with 10 wells of 30 μl. Gels were run for 30 minutes at 200 V, washed with Milli-Q water, stained overnight with InstantBlue and washed three times with tap water. Band intensities were quantified using ImageJ^57^. Background correction was applied to all band intensities by subtracting the average intensity of a region adjacent to the band of interest.

### Fluorescence Recovery After Photobleaching

The bond lifetime of bound crosslinkers was measured via Fluorescence Recovery After Photobleaching (FRAP) using a Nikon A1 confocal microscope with a perfect focus system, 100x 1.40 NA oil immersion objective and 100-mW 488 nm argon ion laser. We acquired 10 images to determine baseline fluorescence and then performed photobleaching by increasing the laser power such that 50-70% of the fluorescence intensity was bleached in 0.5 seconds. We then tracked the fluorescence recovery with a low-intensity beam during a period of approximately 5 times the typical recovery time, with a sampling rate that halved every 10 frames, starting with 10 frames/second. During imaging, the exposure time was kept fixed at 0.1 second/frame. We bleached a circular area of 2 μm radius and used an equally sized area as a reference. The laser intensity during imaging was chosen such that the reference intensity dropped less than 5% during the recovery phase. To extract a timescale for fluorescence recovery, τ_FRAP_, the time-dependent intensity normalized by the intensity of the reference area was fitted with a single exponential function: 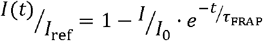, where *I*_0_ is the intensity directly after bleaching^34^.

### Co-sedimentation assay

A volume of 25 μl monomeric (G-)actin at increasing concentrations was co-polymerized with either α-actinin-4 or K255E in F-buffer at room temperature, keeping the crosslinker concentration constant (0.1 μM). After two hours of polymerization, the actin network together with the bound crosslinkers was spun down at 120 000 g. Afterwards, 20 μl was gently pipetted from the supernatant and run on an SDS-PAGE gel as described above. The fraction of bound linkers φ_bound_ was determined by subtracting and normalizing the crosslinker band intensity *I* at a particular actin concentration by the band intensity *I*_0_ in the absence of actin using ImageJ: 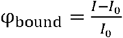.

### Rheology

Rheology was performed using a stress-controlled Kinexus Malvern Pro rheometer with a stainless-steel cone-plate geometry having a radius of 20 mm and a 1-degree cone angle. We loaded 40 μl samples of actin monomers, directly after mixing with either α-actinin-4 or K255E and F-buffer, onto the bottom plate and quickly lowered the cone. A thin layer of Fluka mineral oil Type A was added around the edge to prevent solvent evaporation, and the sample was closed off with a hood to prevent any effects of air flow. Actin polymerization was followed by applying a small oscillatory shear with a strain amplitude of 0.5% and a frequency of 0.5 Hz. After 2 h of polymerization, the elastic shear modulus G’ and viscous shear modulus G’’ were measured as a function of frequency by performing small amplitude oscillatory shear measurements at frequencies between 0.01-10 Hz, taking 30 logarithmically spaced data points. Frequencies above 10 Hz could not be accessed as inertial effects from the rheometer started to dominate the rheological response of the actin network. We used an analytical biopolymer network model to analyze the frequency-dependent viscous shear modulus, *G*’’(*ω*), to extract the crosslinker lifetime in the absence of stress^34,58^. The model is based on the force extension curve of a semiflexible filament, and uses mean-field arguments to calculate the mechanical properties of the network from the single filament fluctuations:

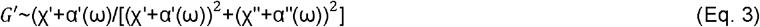

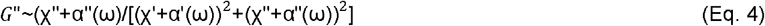

Here, χ describes the viscous drag limiting transverse filament fluctuations:

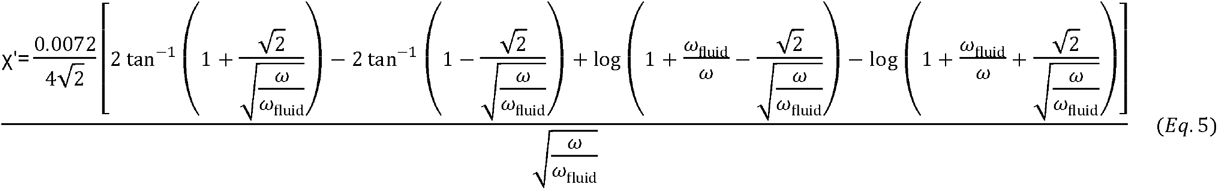

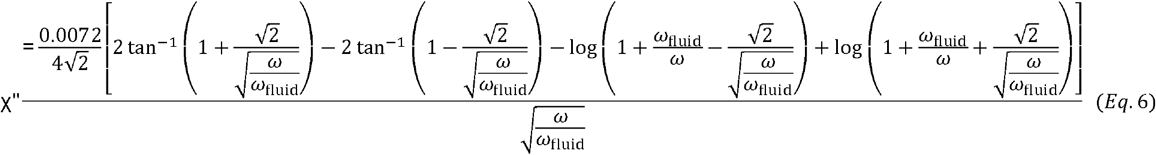

Here ω_fluid_ is the timescale of the fluid drag, which is typically on the order of 100 Hz for actin networks, depending on the fluid viscosity and on the crosslinker and actin concentrations^59^, while α describes the effect of the crosslinkers limiting transverse filament fluctuations:

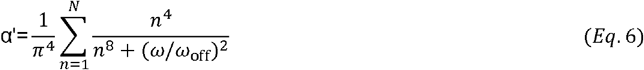

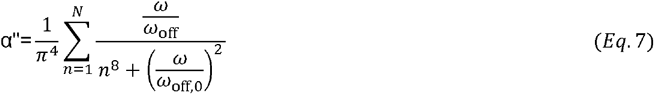

Here *N* is the number of crosslinkers per filament and *ω*_off,0_ is the off-rate of the crosslinker in the absence of force, which we can numerically extract by fitting G”(*ω*) to eqs. 4-7 (Extended Data Fig. 4d). Having characterized the linear rheology, we finally performed a rupture experiment by linearly increasing the stress in time at a constant loading rate (2 mPa/s) until the network ruptured. To unambiguously identify the rupture point, we measured the differential elastic modulus *K’* of the network as a function of stress by superposing small stress oscillations on top of the stress ramp. The rate of 2 mPa/s was chosen because it was sufficiently slow to reliably measure *K’* at every stress level, whilst it was sufficiently fast to prevent network aging effects during the stress ramp. We observed stress-stiffening above a threshold stress, consistent with prior literature^36^, followed by a rapid drop of *K’* that signals rupture. We defined the rupture point as the stress value where *K’* peaked (Extended Data Fig. 4e). This approach allowed us to simultaneously identify the rupture stress and rupture strain (Fig 2c).

To obtain the bond lifetime in the presence of stress, we use a recent extension^36^ of the biopolymer network model described above that takes into account stress-induced network stiffening. Briefly, we superposed an oscillatory stress on top of a constant mechanical load to measure the differential storage modulus *K*’(σ, ω) over a wide range of frequencies (0.01< ω <10 Hz) and stresses (0.1 < σ < 8 Pa) and next fitted the data to the following equation:

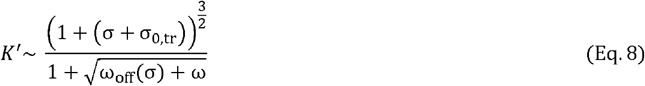

where ω_off_,(σ) is the stress-dependent crosslinker unbinding frequency and σ_0,tr_ is the critical stress for stress-stiffening in the fast limit where crosslinkers have not had time to unbind (Extended Data Fig. 4f-i).

To test whether wall slip contributes to fracturing, we performed control experiments where we fractured actin networks in the absence or presence of a Polylysine-coated glass plate adhered to the bottom and top plates of the rheometer. Polylysine is known to strongly bind actin networks, but neither the rupture strain nor the rupture stress significantly changed upon this surface modification (Extended Data Fig. 10) – showing that fracturing occurred within the network and was not due to wall slip.

### Generation of single-molecule constructs

Both wild-type α-actinin-4 and the K255E mutant were modified to include a ybbR tag (DSLEFIASKLA)^60^right after the His_6_-tag. Purified proteins were coupled to Coenzyme A-modified DNA oligonucleotides 20 nucleotides long using a phosphopantetheinyl transferase (SFP synthase)-mediated reaction^60^. A protein-to-DNA molar ratio of 10:1 ensured that only one monomer was coupled to DNA, as evidenced by SDS-PAGE analysis (Extended Data Fig. 3a). Next, 2.5 kilo base pair DNA tethers were PCR-amplified from the pUC19 plasmid (New England Biolabs) with a 5’-biotinylated primer on one side and a 5’-phosphoprimer on the other side. Purification was done with the QIAquick PCR purification kit (Qiagen, Hilden, Germany). The phosphorylated strand was digested using λ-exonuclease (New England Biolabs) for 2 hours at 37°C and purified using an Amicon 30 kDa MWCO filter (Merck, Darmstadt, Germany). Deep Vent exo-DNA polymerase (New England Biolabs) and a 20-nucleotides more upstream primer than the phosphoprimer from the PCR was used to fill up the second DNA strand, creating a 20-nucleotide overhang^61^. This overhang is complementary to the 20-oligonucleotide sequence coupled to the proteins. The generated DNA tether was then ligated to the DNA-protein hybrid by overnight incubation with T4 ligase (New England Biolabs) at room temperature. The stock sample was flash frozen and stored at -80 °C, and small aliquots were stored at 4 ^º^C for maximally one week.

### Preparation of actin-coated and crosslinker-coated beads

Unlabeled actin monomers were mixed with biotinylated monomers and fluorescent monomers labeled with Alexa Fluor 647 in a molar ratio of 8:1:1 and polymerized into filaments in 1 mL F-buffer at a concentration of 2 μM for 2 hours. Next, these filaments were mixed with 4 μl of 2.4 μM Neutravidin-coated beads (NVP-20-5, diameter 2.1 μm, Spherotech) and incubated for 15 minutes to couple the filaments to the beads. The actin-coated beads were separated from unbound actin filaments by centrifuging 3x at 1000 rcf for 2 minutes. After every round, 800 μl of supernatant was discarded, whilst carefully avoiding disturbing the pellet, and replaced by 800 μl of fresh F-buffer. Successful coating was verified using confocal fluorescence microscopy by the presence of a fluorescent ring on the edge of the bead upon excitation with a 638 nm laser (Extended Data Fig. 3b). For the other bead type, approximately 50 ng of the generated crosslinker-DNA construct was incubated with 2 μl NeutrAvidin beads in 10 μl F-buffer for 15 min in a rotary mixer at 4□°C, and then rediluted in μl 500 F-buffer with 100 mM biotin excess to block unbound NeutrAvidin. Unbound biotin was removed during the optical tweezer assay by flushing F-buffer after trapping the beads.

### Single molecule data acquisition and analysis

Force spectroscopy data was collected at 500 Hz using a custom-built dual trap optical tweezers and a commercial C-Trap (Lumicks). Data was analyzed using custom scripts in Python. The optical traps were calibrated using the power spectrum of the Brownian motion of the trapped beads^62^, obtaining average stiffness values of κ = 0.39 ± 0.04 pN nm^-1^. After trapping beads with the two different coatings (Extended Data Fig. 3b), α-actinin-4-actin binding was established by approaching and maintaining both beads in close proximity during approximately 10 seconds. Tether lifetime was assessed by rapidly retracting the beads to a set distance – thus increasing the applied force – and measuring the time until the tether broke. To discriminate single from multiple connections, we used the worm-like-chain (WLC) model and the fact that single double-stranded DNA exhibits an overstretching plateau in the force-extension curve at forces above 65 pN (Extended Data Fig. 3c). We pulled on tethers to high forces and observed that the contour length (computed using the WLC) of those that displayed overstretching characteristic of single tethers matched the expected value of 850 nm within a ∼60 nm range, likely due to the variability in the bead radii and the thickness of the actin coat. Multiple tethers, in contrast to single ones, did not show this characteristic overstretching, and their apparent length was most often shorter (Extended Data Fig. 3c). Therefore, we considered tethers that displayed the expected contour length of 850 ± 30 nm and broke in a clean step. Most tethers showed dissociation below a minute waiting time (55% of tethers, across all forces). Tethers that lasted longer generally did not break at all, even after several minutes under tension. Hence, lifetimes were determined from tethers showing dissociation within one minute. Numbers of data points per bin in Fig. 1f are: 6, 5, 7, 2, 4, 2, 2, 1, 0 for WT, and 3, 7, 7, 4, 4, 8, 5, 5 for the K255E mutant.

## Author contributions

Y.M. and G.H.K. conceived and designed the study. M.J.A. and S.J.T. designed the optical tweezer experiments. M.J.A., A.R. and L.B. performed the optical tweezer experiments. M.J.A. and A.R. analyzed the optical tweezer experiments. Y.M. performed and analyzed all other experiments and designed and simulated the 1D model. W.J. and T.K designed, performed and analyzed the actin network simulations. Y.M., M.J.A, T.K. S.J.T. and G.H.K. wrote the manuscript with input from W.J., A.R. and L.B. All authors approved the final version.

## Acknowledgements

We thank Martin van Hecke and Celine Alkemade for critical reading of the manuscript. We thank Pieter Rein ten Wolde, Kees Storm, Wouter Ellenbroek, Chase Broedersz, David Brueckner and Mareike Berger for fruitful discussions. We thank William Brieher and Vivian Tang for the kind gift of purified α-actinin-4 (wild type and the K255E point mutant) and their plasmids, Marjolein Kuit-Vinkenoog and Jeffrey den Haan for actin and further purification of α-actinin-4, and Vanda Sunderlíková for design, mutagenesis, cloning and purifying of the α-actinin-4 constructs used in the single molecule experiments. This work is part of the research program of the Netherlands Organization for Scientific Research (NWO). We gratefully acknowledge financial support from an ERC Starting Grant (335672-MINICELL) awarded to G.K and by the “BaSyC – Building a Synthetic Cell” Gravitation grant (024.003.019) of the Netherlands Ministry of Education, Culture and Science (OCW) and the Netherlands Organisation for Scientific Research. We gratefully acknowledge the support from the National Institutes of Health (1R01GM126256).”

## Data availability

The data that support the findings of this study are available from the corresponding authors upon reasonable request.

## Code availability

Custom-written scripts used in this study are available from the corresponding authors upon reasonable request.

**Supplementary Information** is available for this paper.

**Supplementary Video 1** Visualization of the top 10% stressed filaments in actin network simulations of catch (top) and slip bond networks (bottom) with actin filaments in green and the network borders in white. The network is strained over time by moving the top boundary. The x- and y-axis shows the distance in μm.

**Supplementary Video 2** This video shows the same analysis as Fig. 3e, with each frame showing the distribution at a different stress level (top right corner). Color scale: percentage relative enrichment of catch bonds over slip bonds bound between a pair of actin filaments for varying tension on filament 1 (vertical axis) and on filament 2 (horizontal axis). Specifically, the color scale is defined as 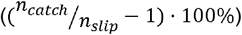 when *n*_*catch*_ > *n*_*slip*_ (shown in red) and 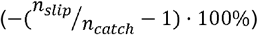 when *n*_*slip*_ > *n*_*catch*_ (shown in blue) where *n*_*catch*_ and *n*_*slip*_ are the number of resp. catch and slip bond linkers connecting two filaments. The tension spacing is the same for the x- and y-axis and is updated at every timestep such that each bin includes 10% of the filaments with the top-right corner presenting connections between the tensest filaments.

**Correspondence and requests for materials** should be addressed to G.H.K, S.J.T and T.K.

## Supplementary Information

### Experiments

#### Frequency-dependent rheology shows that crosslinking affects the dynamics, but not the structure of actin networks

We studied the influence of the bond lifetime of the two α-actinin-4 variants (wild type and K255E) on the dynamics of actin networks using small-amplitude oscillatory shear measurements on the crosslinked actin networks over a range of oscillation frequencies (10^−3^ to 10 Hz, see Extended Data Fig. 4a). For both variants, the frequency spectrum of the shear moduli has a functional form that is characteristic of transiently crosslinked semiflexible polymer networks^1,2^. For low frequencies, below the crosslinker unbinding frequency (inverse bond lifetime), both networks deform viscoelastically, with the storage modulus *G*’ and loss modulus *G*’’ both following a power law dependence on frequency with an exponent of 1/2. This exponent arises from the superposition of multiple relaxation times of many crosslinkers connecting a single filament to the surrounding network^1^. We observe a characteristic relaxation frequency where *G*’’ exhibits a peak and *G*’ exhibits an elastic plateau (Extended Data Fig. 4a-c). This relaxation frequency corresponds to the crosslinker unbinding rate^1^. At frequencies above 5 Hz, both moduli show a slight upturn, which reflects the influence of viscous drag on the actin filaments.

When we compare the frequency spectrum for actin networks crosslinked with the two α-actinin-4 variants, we observe that replacing the wild type variant by the K255E mutant causes a shift to a lower relaxation frequency, indicating slower unbinding as expected from the higher binding affinity. When we normalize the applied frequency with the relaxation frequency, the *G’* and *G”* curves for the K255E mutant can be super-imposed on the wild type α-actinin-4 curves with only a small deviation at high normalized frequencies, which we attribute to the effect of the viscous drag on the filaments (Extended Data Fig. 4b). This collapse suggests that, although the crosslinker lifetime is shorter for the wild type α-actinin-4 than for the K255E mutant, the network structure is not significantly different. For example, any change in the typical crosslinker distance would have altered the shear modulus^1^. Indeed, with light microscopy we do not find any bundles, indicating that both networks are isotropically crosslinked (Extended Data Fig. 9).

We tune the relaxation frequency for actin networks crosslinked with K255E by increasing the temperature^3^ from 10 ^º^C to 25 ^º^C such that it equals the peak frequency of networks crosslinked with the wild type variant at low temperature (10 ^º^C). Now, the time-dependent linear rheology of both networks is indistinguishable even without normalizing the frequency (Extended Data Fig. 4c-d).

### Minimal crosslinker model

#### Discussion of the minimal crosslinker model design

The aim of our minimal 1D model is to identify the essential ingredients necessary to explain how catch bonds can make stronger networks despite being weaker on the single bond level. Therefore, the model focuses on the key difference of networks crosslinked with wild type (catch bond) and mutant (slip bond) α-actinin-4 crosslinkers, namely the force-dependent bond dynamics (Methods Eq. 1). The only parameter we vary between catch and slip bonds is 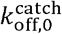 (0 for slip bonds and positive for catch bonds). This parameter represents the activation of a binding site as a function of force, with 0 meaning constitutive exposure. The resultant theoretical force-lifetime curve (Fig. 1b) is qualitatively consistent with the single molecule force spectroscopy data and with what is known about the structure of α-actinin-4, specifically the hidden ABS1-domain (Fig. 1a, e, see main text Introduction)^4,5^. Apart from the bond dynamics, we need the minimal ingredients to allow for crack initiation and propagation in our model. This requires spatial inhomogeneity, which is already possible in one dimension. We implement this inhomogeneity in the simplest possible form, namely by nearest-neighbour load sharing (Methods Eq. 2), which suffices to study crack propagation - as we have previously shown and benchmarked with finite element simulations^6^. These minimal ingredients (force-dependent bond dynamics and nearest-neighbour load sharing) not only suffice as a minimal model for crack initiation in viscoelastic materials^6^, but also qualitatively reproduce the key result of our studies, namely that weak catch bonds make strong networks (Fig. 2b). Further, it correctly predicts that the majority of crosslinkers is bound (Extended Data Fig. 1), and that the rupture stress increases more strongly with bond lifetime for catch bonds than for slip bonds (Extended Data Fig. 7).

The simplicity of the model allows for a conceptual interpretation of the experimental findings and identifies the ingredients that are necessary and sufficient, i.e., a large system, bond mobility, and force-activated binding. However, the lack of detail in the model regarding the actual 3D network architecture does put limits to its applicability. First, the model contains crosslinker inhomogeneity but lacks the *network* inhomogeneity inherent to any realistic biopolymer network such as inhomogeneous polymer distribution and stress-induced alignment. Network inhomogeneity is known to strongly enhance fracturing^7^. However, other factors reduce the likelihood of fracturing such as force correlations between crosslinkers in real networks are likely more long-ranged than only nearest-neighbor load sharing. Therefore, one should not expect quantitative predictions of rupture stresses and rupture strains from the model. We expect that including network inhomogeneity will only further enhance the strength advantage of catch bonds over slip bonds, as catch bonds self-assemble against the force inhomogeneity induced by the network inhomogeneity, consistent with the bigger difference in rupture stress in the experiments (>3x increase in the experiments vs. a ∼2x increase in the minimal model).

Second, as the model is strictly speaking kinetic rather than mechanical, it does not explicitly model deformations. We use bond turnover as a proxy for deformability. Indeed, actin networks without bond turnover rupture at only approximately 10% shear strain^8^, whereas the transiently crosslinked networks here reach maximal strains of up to several hundred percent, showing that bond turnover dominates network deformability. However, the exact, quantitative relationship is unknown^9,10^ and beyond the scope of this work. As a result, only qualitative comparisons on the deformability between real and simulated networks can be made (Fig. 2b,d, y-axes). With this caveat, the network does correctly reproduce the qualitative feature that the network dynamics increases with decreased crosslinker bond lifetime and with catch bonding. In contrast, our detailed actin network simulations do show rupture strains which can directly be compared to the experiments and show quantitative agreement between simulated and experimentally measured rupture strains within a factor of 2. The actin network simulations likely somewhat underestimate the rupture strain as the simulated networks are deformed at a much higher rate than in the experiments for computational reasons

Thirdly, the relevance of our model is limited to crack initiation, and crack propagation is out of the scope of this model as the fracture energy during crack propagation depends on network-specific physical processes, such as solvent dynamics and polymer disentanglement. Since crack initiation is the rate-limiting step in fracturing under constant stress, the model suffices for our application but might be insufficient in addressing fracturing during other modes of deformation.

#### Quantitative comparison between the minimal model and the different experiments

Although the 1D model is primarily designed to qualitatively study the mechanism of fracturing, as detailed above, a rough quantitative comparison with the force/stress levels in the experiments can nevertheless be made. Here we show how to extract the average force per crosslinker in the rheology experiments, which we then combine with the single molecule force spectroscopy findings to enable a direct comparison with the model. We perform our calculations on the networks at room temperature (300 K), as this is the only condition where we have both network rheology and single molecule force spectroscopy data.

In order to obtain the average crosslinker force from the network stress, we need the typical crosslinker distance, *l*_*c*_, which can be computed from the measured network elasticity G_0_ using the following expression, derived for isotropic semiflexible polymer networks^11^:

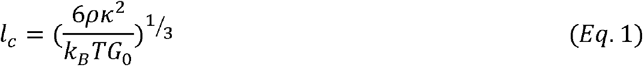

Here, *κ* is the filament bending modulus (7.14·10^−26^ J·M for actin^12^), *k*_*B*_ is the Boltzmann constant, *T* is the temperature in *K, G*_0_ is the plateau elastic modulus (around 200 Pa in all of our networks, Extended Data Fig. 4 a-c) and *ρ* is the filament line density (in m^-2^), which can be calculated from the actin concentration, *c*_a_ (in units of mg/mL) following simple geometry:

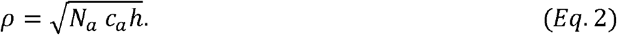

Here, *N*_a_ is the Avogadro constant and *h* the length each actin monomer adds to the polymer (roughly 2 nm, as the polymer is a double helix and monomer diameter is roughly 4 nm^12^). As we have used 48 μM actin in our experiments, the line density is 6.72·10^13^ m^-2^. Lastly, we calculate the force per crosslinker *f*_*c*_ from the network stress σ, by assuming the stress is homogeneously distributed over meshes consisting of four crosslinkers each^13^:

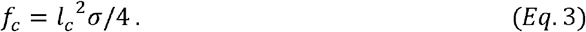

As a result, we find that the average crosslinker force in our experiments is around 1.8 pN per 1 Pa of applied network stress. Therefore, the average crosslinker force at rupture is about 11 pN for slip bonds and about 13 pN for catch bonds. These forces are in the same range as the force range where we observe catch bonding for the wild type crosslinkers in the single molecule force spectroscopy. An independent confirmation that the force ranges in the rheology and single molecule experiments agree and that catch bonding occurs in both cases is that frequency-dependent rheology measurements reveal that the wild type crosslinker shows a significant increase of the stress relaxation time with increasing applied stress, whereas the mutant shows a continuous decrease (Extended Data Fig. 4f-h).

Before we can compare the experimentally applied network stress to the normalized network stress in the simulations, we need to extract the crosslinker force sensitivity from the single molecule force spectroscopy (Fig. 1f). In the model, the force sensitivity is defined as the force at which the crosslinker lifetime is reduced by a factor *e* (see Eq. 1 in the Methods section for more details). Fig. 1f shows that experimentally, this occurs roughly at 10 pN. We can use this value to directly compare the normalized stress to the stress applied by rheology and find that a normalized stress of 1 is approximately equal to 5.5 Pa. The model therefore predicts rupture stresses on the order of a few Pascal, consistent with the magnitude of the rupture stresses observed in the experiments. Lastly, we note that the force sensitivity of the actin-actin interface within an actin polymer is ∼50 pN^14^, making the crosslinker-actin interface by far the weakest link of the network and therefore the point of fracturing.

#### Catch bonds still enhance network strength when taking into account their dimeric molecular structure

In the minimal 1D crosslinker model presented in the main text, crosslinkers are either bound or unbound. However, the α-actinin-4 crosslinkers in the experiments and in the polymer network simulations can also be partially bound (i.e., only to one actin filament), because they have two independent actin-binding domains separated by a spacer. To test whether allowing for partially bound crosslinkers qualitatively affects the model predictions, we expand the model to allow for three binding states, where doubly bound crosslinkers become singly bound crosslinkers with an unbinding rate of *k*_2→1_(*f*_*i*_), following force-induced unbinding according to Eq. (1) of the Material and Methods. Singly bound crosslinkers become fully unbound with a rate of *k*_1→0_ or doubly bound with a rate of *k*_1→2_ Force is only shared between neighboring doubly-bound crosslinkers. For simplicity, we have chosen *K*_1→0_ = *k*_2→1_ (0) and *k*_0→1_ *= k*_1→2_ = 1. Importantly, allowing for this additional, partially bound state does not qualitatively affect the predictions of the computational model, as catch bonds still form networks that are both stronger and more dynamic than networks formed with slip bonds (Extended Data Fig. 6d). For all other simulations, we therefore use a two-state model as it is the most minimal model that captures our experimental results.

#### Catch bonds enhance the strength of materials that fracture via crack propagation

Our experiments demonstrate that catch bonds collectively provide stronger networks, even though they are actually weaker on the single molecule level. This raises the question how many bonds are required for the catch bond advantage to emerge. To answer this question, we performed fracturing simulations at different network sizes (1-200 linkers, Extended Data Figure 7a,b). We find that for both catch and slip bond networks, the rupture strength and bond turnover number initially increase with increasing bond number, peak when the bond number reaches around 10, and then continuously decrease with increasing bond number. Remarkably, slip bond networks are stronger for systems smaller than 10 bonds, while catch bond networks have both larger network strength and bond turnover for networks larger than 10 bonds.

Why is catch bonding (only) superior in large networks? To understand this, we need the notion of a critical crack size from fracture mechanics^6,15^. When a large crack (a part of the network that is devoid of crosslinkers) is under stress, bonds at the edge of the crack rapidly unbind, causing crack propagation and eventually network fracturing. However, when the crack is still small, linker binding may heal the crack before it spreads. Therefore, there is a critical crack size that is on the verge of becoming unstable. For systems below this critical crack size, the network strength increases with the bond number as it becomes increasingly unlikely that all linkers simultaneously unbind. For large systems however, increasing the network size allows for more locations at which a crack can be initiated – causing a decrease in network strength. Combined, these two effects explain why we observe a biphasic dependence of network strength on system size, with a cross-over at a critical crack size (Extended Data Fig. 6a,b, see Refs.^6,16^ for a more detailed and quantitative explanation).

So why are catch bonds only effective for systems larger than the critical crack size? Catch bonds rely on bond redistribution to enhance the network strength (Extended Data Fig. 6c). The ‘dissociation-on-demand’ of unforced catch bonds increases the pool of unbound crosslinkers and thereby crosslinker binding in the entire network, causing a net crosslinker migration from stress-free to high-stress areas, as can be seen from the narrow force distribution (Fig. 3a). This increased binding in high-stress areas makes cracks less prone to becoming unstable (Fig. 3b), thus increasing the network strength. However, this effect relies on catch bonds to migrate from outside of the critical crack size towards the crack, and therefore only emerges in networks larger than the critical crack size.

Although it is difficult to precisely know the critical crack size for viscoelastic materials, any macroscopic network that can be studied with bulk rheology techniques is orders of magnitude above this size threshold^6^. Furthermore, laser ablation experiments have revealed that cells are also significantly larger than the critical crack size^17^. Therefore, we conclude that the catch bond advantage is general and should emerge both in cells and macroscopic networks. Lastly, as cell adhesion typically also relies on hundreds of linkers^18^, catch bonds likely also strengthen cell-cell and cell-matrix adhesions whilst simultaneously facilitating sliding. Therefore, our results could also explain why biological adhesins are very often catch bonds^19^.

### Actin network simulations

#### Simplification of actin filaments (F-actin) and crosslinkers

Network simulations were implemented using procedures described in detail in earlier works^20–23^. We provide a brief summary here. F-actin and crosslinker are simplified using cylindrical segments (Fig. 3c). F-actin is simplified into cylindrical segments of 420 nm in length with polarity (i.e., barbed and pointed ends) that are serially connected via elastic hinges. Each crosslinker is comprised of two arms connected at its center point via elastic hinges. Binding sites for the arms of crosslinkers are located on each actin segment every 7 nm.

#### Brownian dynamics via the Langevin equation

The displacement of each cylindrical segment constituting F-actin and crosslinker is regulated via the Langevin equation with inertia neglected:

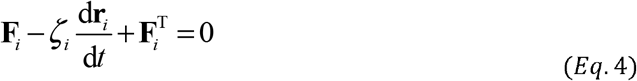

where *F*_*i*_ is a net deterministic force, *ζ*_*i*_ is a drag coefficient, *r*_*i*_ is a position, *t* is time, and *F*_*i*_^T^ is a stochastic force satisfying the fluctuation-dissipation theorem^24^:

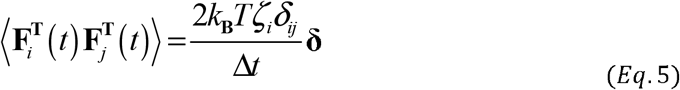

where *δ*_*ij*_ is the Kronecker delta, Δ*t* = 5.94×10^−5^ s is a time step, and **δ** is a unit second-order tensor. *ζ*_*i*_ is calculated by the approximate equation for a cylindrical object^25^:

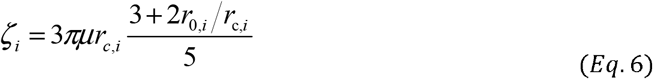

where *μ* is the viscosity of the surrounding medium, and *r*_c,*i*_ and *r*_0,*i*_ are the diameter and length of a segment, respectively. At each time step, the position of each segment is updated via the Euler integration scheme:

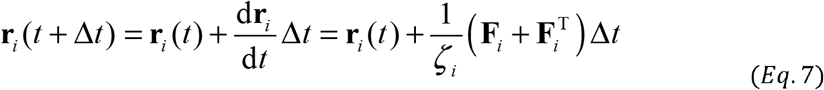

#### Deterministic forces

The deterministic force, *F*_*i*_, includes extensional, bending, and repulsive forces. The extensional and bending forces are calculated based on harmonic potentials:

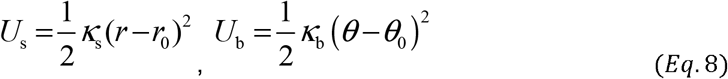

where *κ*_s_ and *κ*_b_ are extensional and bending stiffnesses, respectively, *r* is the length of a segment, θ is an angle formed by segments, and the subscript 0 represents an equilibrium value. The extensional (*κ*_s,A_) and bending (*κ*_b,A_) stiffnesses of F-actin maintain the length of each actin segment at an equilibrium length (*r*_0,A_ = 420 nm) and an angle formed by adjacent actin segments at an equilibrium angle (*θ*_0,A_ = 0°), respectively. Similarly, the extensional (*κ*_s,x_) and bending (*κ*_b,x_) stiffnesses of crosslinkers maintain the length of each crosslinker arm and an angle formed by two crosslinker arms at equilibrium values (*r*_0,x_ = 23.5 nm and *θ*_0,x_ = 0°), respectively. Forces exerted on the binding sites of an actin segment by crosslinkers are distributed to two endpoints of the segment as explained in our previous work^21^.

Repulsive forces between neighboring actin segments are calculated based on the following harmonic potential:

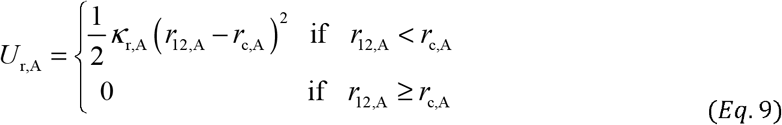

where *r*_12,A_ is a minimum distance between two neighboring actin segments, and *κ*_r,A_ is strength of the repulsive force.

#### Crosslinker dynamics

Crosslinkers exist in one of the three states: monomeric, inactive, and active states. Monomeric crosslinkers that are not bound to any F-actin are considered implicitly by their local concentrations. The monomeric crosslinkers bind to binding sites located on actin segments at a rate calculated using a binding rate, *k*_+,x_, and their local concentration. Then, they become inactive crosslinkers that are bound to only one F-actin. The inactive crosslinkers bind to binding sites located on another actin segment without any preference for a crosslinking angle at the constant binding rate, *k*_+,x_. For the binding event, a distance between the actin segment and the center point of the inactive crosslinker should be between 21.2 nm and 25.9 nm (± 10% with respect to *r*_0,x_). After the binding event, they become active crosslinkers.

The active crosslinkers also unbind from F-actin in a force-dependent manner, following Bell’s law:

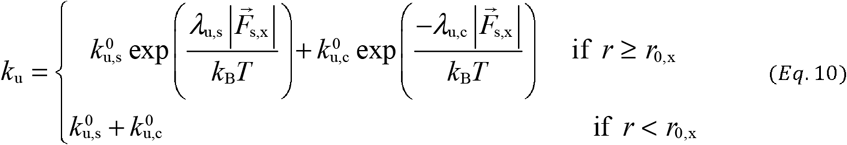

where *k*^0^_u,s_ and *k*^0^_u,c_ are zero-force unbinding rate constants for slip and catch bond parts, respectively, and *λ*_u,s_ and *λ*_u,c_ represent sensitivity to the magnitude of applied spring force 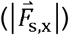 for slip and catch bond parts, respectively, and *k*_B_*T* is thermal energy. Note that we set *k*^0^_u,c_ to zero or a finite value to simulate slip or catch bonds.

#### Network assembly

At the beginning of each simulation, a crosslinked actin network is assembled via dynamic events of F-actins and crosslinkers in a three-dimensional thin rectangular domain (30×30×1 μm) with a periodic boundary condition in x and y directions (Fig. 3d). The nucleation of F-actins occurs in a random direction perpendicular to the z direction, followed by relatively fast polymerization. The nucleation of F-actin is represented by the appearance of one cylindrical segment occurring at a given nucleation rate, *k*_n,A_. The polymerization is represented by the addition of cylindrical segments to the barbed end at a given assembly rate, *k*_+,A_. This results in formation of a network with randomly oriented F-actins. While F-actins are nucleated and polymerized, crosslinkers bind to a pair of F-actins to form a functional crosslink between F-actins.

#### *In silico* bulk rheology

After network formation, F-actins at the upper (+y) and lower (-y) boundaries are severed and clamped, and the periodic boundary condition is removed in the y direction. F-actins do not undergo dynamic events. Then, a shear strain with a constant strain rate, 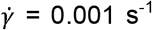, is imposed. Actin segments located within 420 nm from the bottom surface are fixed, whereas those located within 420 nm from the top surface are displaced following linearly increasing shear strain. The resultant stress exerted by the network is measured by summing the x component of the forces acting on the ends of F-actins clamped to the top surface and then dividing the sum by the area of the top surface. Because stress is output from simulations, it is hard to precisely control the stress without a noise in simulations, so we controlled strain as input in simulations rather than stress.

## Extended Data Figures

**Extended Data Fig. 1:**
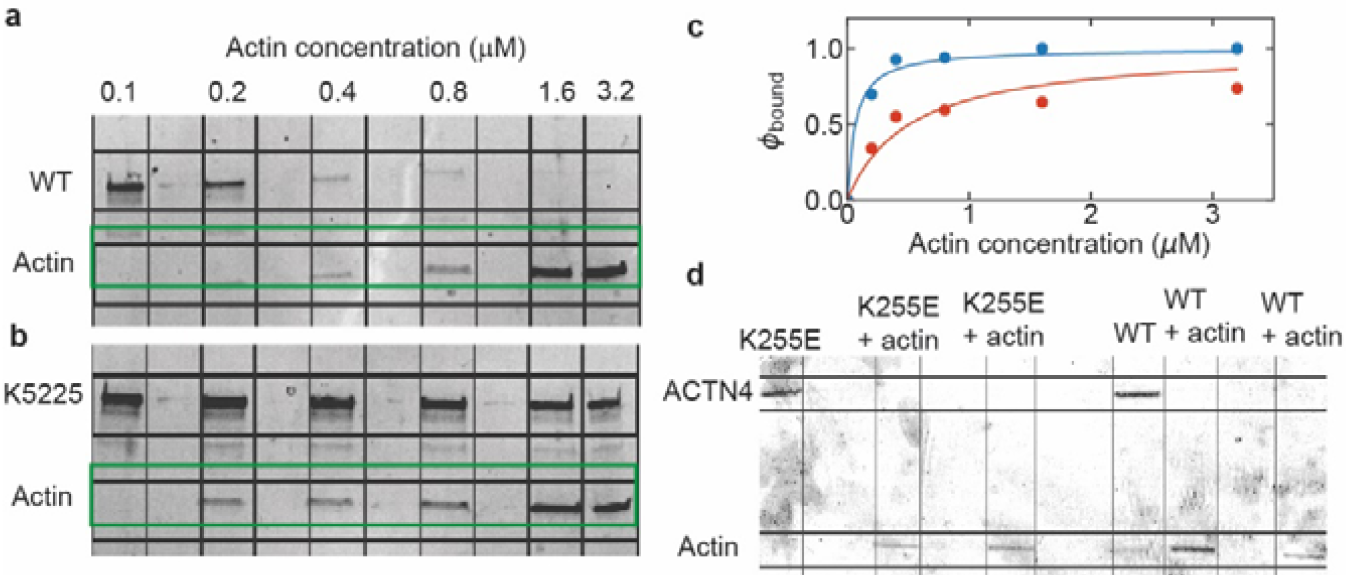
High-speed co-sedimentation measurements of the affinity of α-actinin-4 (WT) and K255E crosslinkers for actin filaments. **a, b**, supernatant resulting from a high-speed centrifugation of a mixture of actin filaments and crosslinkers was run on an SDS-page gel. The bands on the bottom show the α- actinin-4 (WT) or K255E (resp. **a** and **b**, molecular weight ∼ 100 kDa in both cases), while the bands on the top show actin (42 kDa). Each labeled column contained a different actin concentration as indicated. Some lanes were kept empty as spacers. The crosslinker concentration was fixed at 0.1 μM. **c**, The fraction of bound crosslinkers, as determined from the co-sedimentation assay, as a function of the actin concentration was fit to the equation: 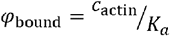, where *K*_*a*_ is the affinity of the crosslinker. **d**, Consistent with the high affinity of both crosslinkers, SDS-page gels of supernatant resulting from a high-speed centrifugation of a crosslinked actin network at the concentration used in all our experiments (48 μM actin together with 0.48 μM crosslinker) does not show any measurable fraction of unbound crosslinkers.

**Extended Data Fig. 2:**
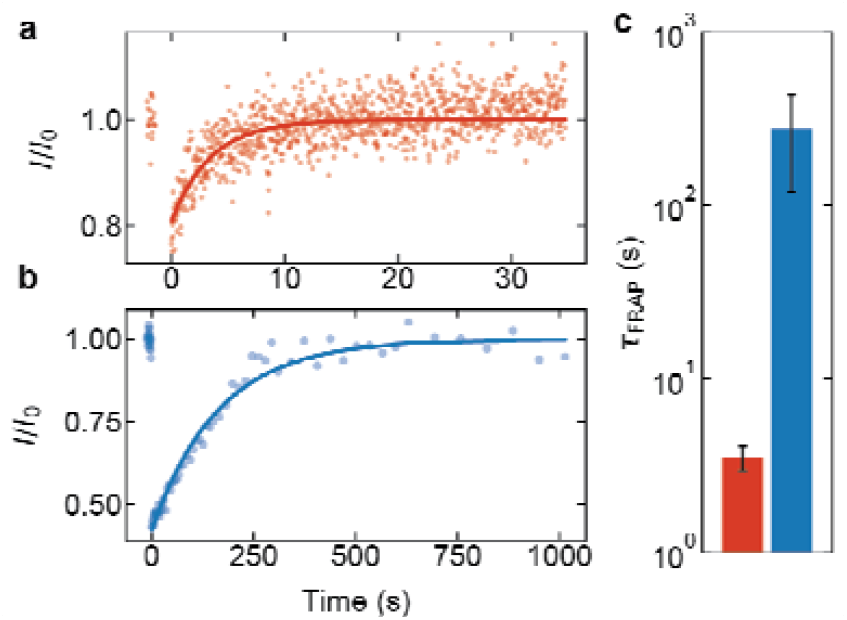
Fluorescence recovery after photobleaching measurements reveal that α-actinin-4 crosslinkers are more dynamic than the K255E mutant. Example fluorescence recovery curves of α-actinin-4 (**a**) and K255E (**b**) in the presence of 48 μM actin show full recovery of both proteins after photobleaching at time *t*=0, but with different timescales. The solid lines represent exponential fits to the data (see Methods). **c**, Average recovery time for α-actinin-4 (red) and for K255E (blue), with the standard error on basis of 6 repeats per condition. Measurements were performed at 25 °C.

**Extended Data Fig. 3:**
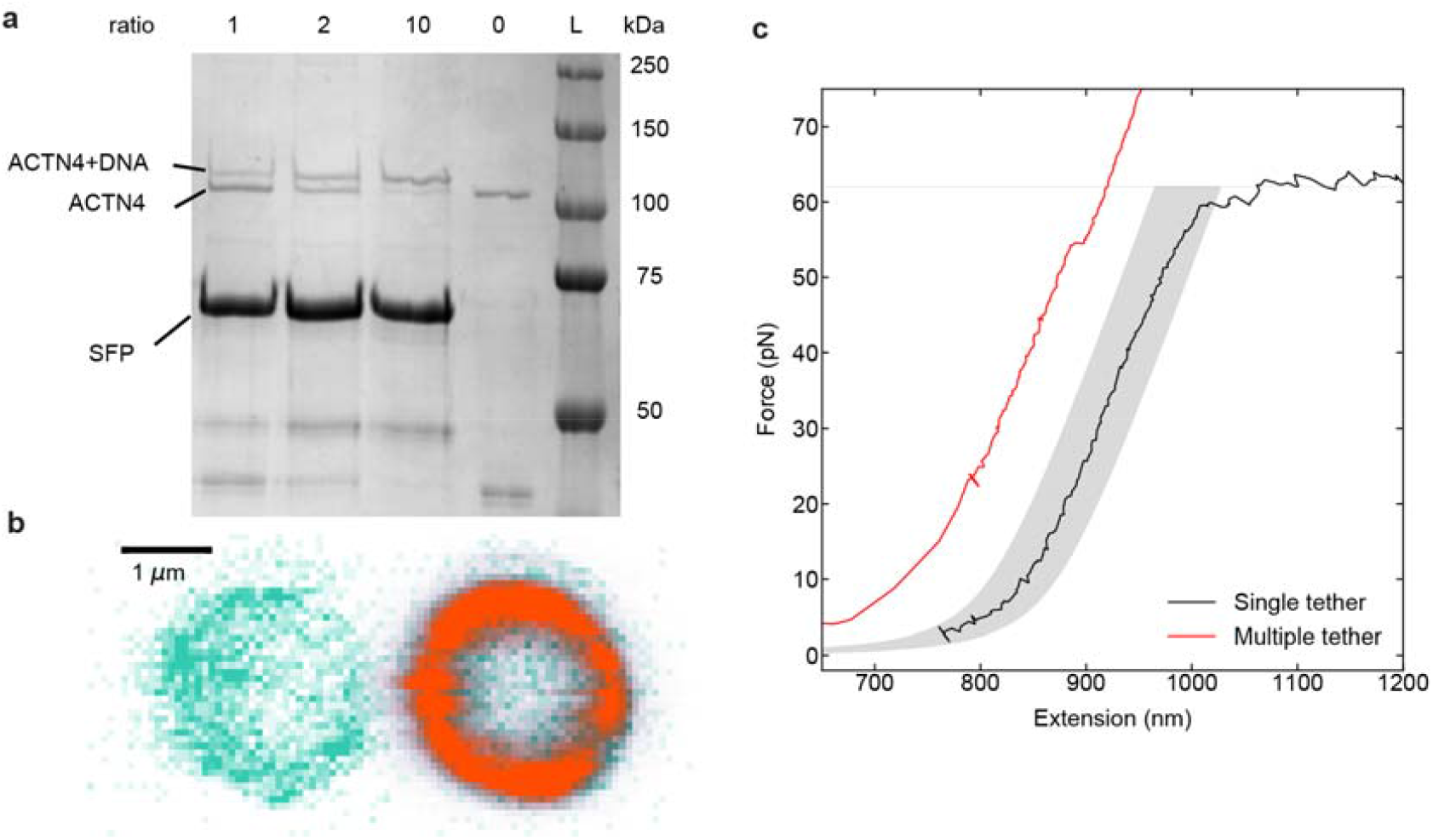
Generation and classification of α-actinin-4/actin tethers. **a**, DNA was coupled to α-actinin-4 (WT or K255E) using an SFP synthase-mediated reaction. Because α-actinin-4 is a homodimer, the yBBr tag used for coupling is present in both monomers. To favour DNA attachment to only one monomer, we performed coupling reactions with several DNA titrations, and the coupling yields were quantified using SDS-PAGE gel electrophoresis. The DNA:α-actinin-4 molar ratios are indicated above each lane. At a molar ratio of 1:1, most of the α-actinin-4 is uncoupled, i.e. most dimers will be either not coupled or have only one monomer coupled to DNA. **b**, Concurrent confocal fluorescence images of a trapped bead coated with α-actinin-4 (left) and a trapped bead coated with actin filaments (right). The bead’s autofluorescence is depicted in green, and the fluorescent emission of Alexa Fluor 647-tagged actin is depicted in orange. **c**, Force-extension curves showing the overstretching regime of a single dsDNA tether (black), and a case where the two beads are linked by multiple tethers, which yields a shorter contour length and higher forces without unzipping (red). Variability in bead radii and actin layer thickness results in force-extension curves that can be shifted along the Extension axis, from the theoretical 850 nm by ±30 nm. Grey area: “single-tether region”. Tethers with a force-extension curve within this area that broke in a single step were regarded as single tethers and hence included in measuring the force-dependent lifetime.

**Extended Data Fig. 4:**
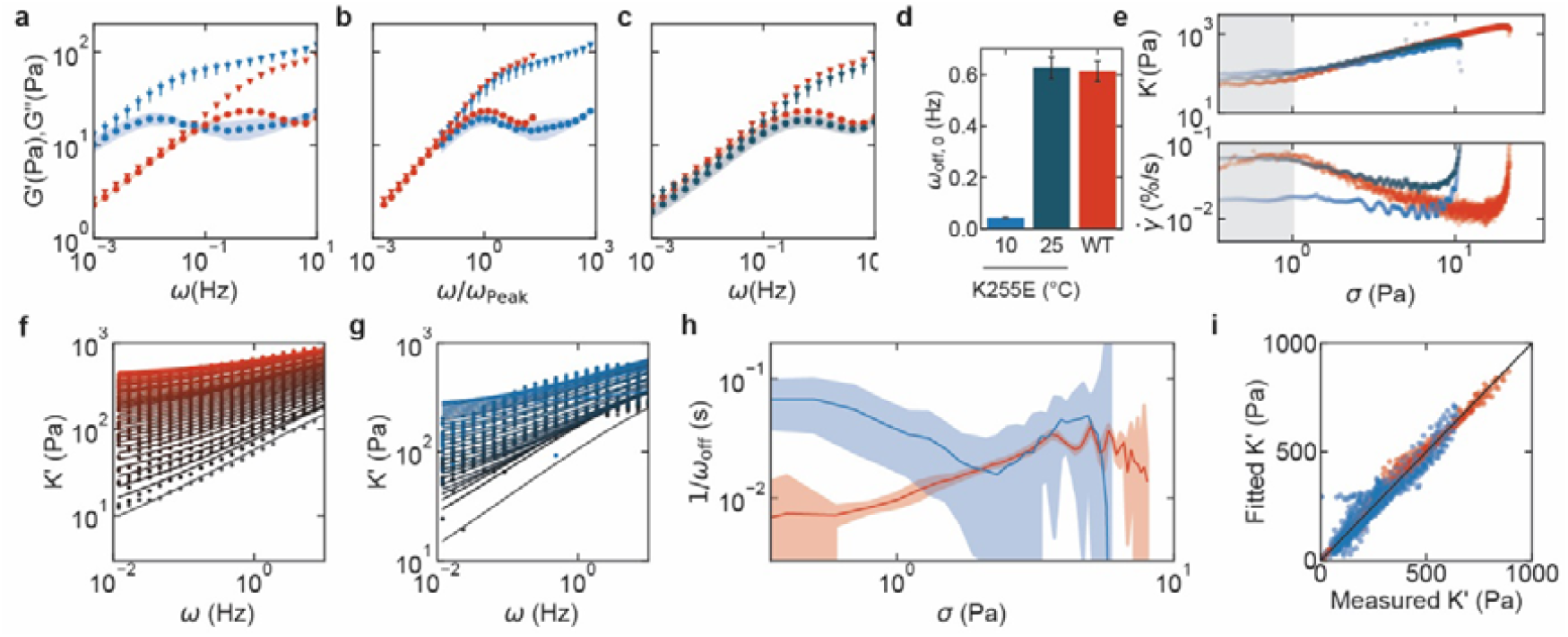
Nonlinear and temperature-dependent rheology of actin networks crosslinked by α-actinin-4 or K255E. **a-c**, The storage (triangles) and loss moduli (circles) were measured using small amplitude oscillatory shear. The moduli are shown as a function of frequency (**a**) and as a function of the frequency normalized by the frequency at which the loss modulus peaks (**b**). The peak frequency is 0.5 Hz for α-actinin-4 (red), and 0.01 Hz for the K255E mutant (light blue). Both curves are measured at 10 °C. **c**, The time-dependent rheology of actin networks is compared between α-actinin-4 crosslinking at 10 °C (red) and K255E crosslinking at 25 °C (dark blue). The standard error indicated by bars and shaded regions is on basis of 4 repeats per condition. The collapses in **b** and **c** show that the crosslinker unbinding kinetics, but not the network structure, is significantly different for the different conditions (see Main Text). **d**, The stress relaxation frequency was extracted from Extended Data Fig. 4a-c using Methods Eq. 4-7 for α-actinin-4 WT at 10 °C (red), K255E at 10 °C (light blue) or K255E at 25 °C (dark blue). **e**, Representative example curves of the differential storage modulus at 0.5 Hz (top) and of the strain rate (bottom) are plotted against the applied shear stress for actin networks crosslinked by α-actinin-4 WT at 10 °C (red), K255E at 10 °C (light blue) or K255E at 25 °C (dark blue). We define the rupture strain as the data point where K’ peaks. **f-g**, We apply a semiflexible polymer network model to fit the frequency-dependent differential elastic modulus as a function of prestress (see Methods). **h**, Thus, we extract the crosslinker bound lifetime as a function of stress for both α-actinin-4 (red) and the K255E mutant (blue) at 25 °C. The shaded areas represent the error on basis of the fits. The bound lifetime of the mutant is significantly longer at low stress, but the lifetimes of catch and slip bonds become similar at high stress as the bound lifetime of the catch bonds increases. The abrupt decay of bound lifetime in the K255E-crosslinked network when the stress reaches 5 Pa is due to network fracturing. **i**, the fitted K’ shows quantitative agreement with the measured K’ for both catch and slip bonds.

**Extended Data Fig. 5:**
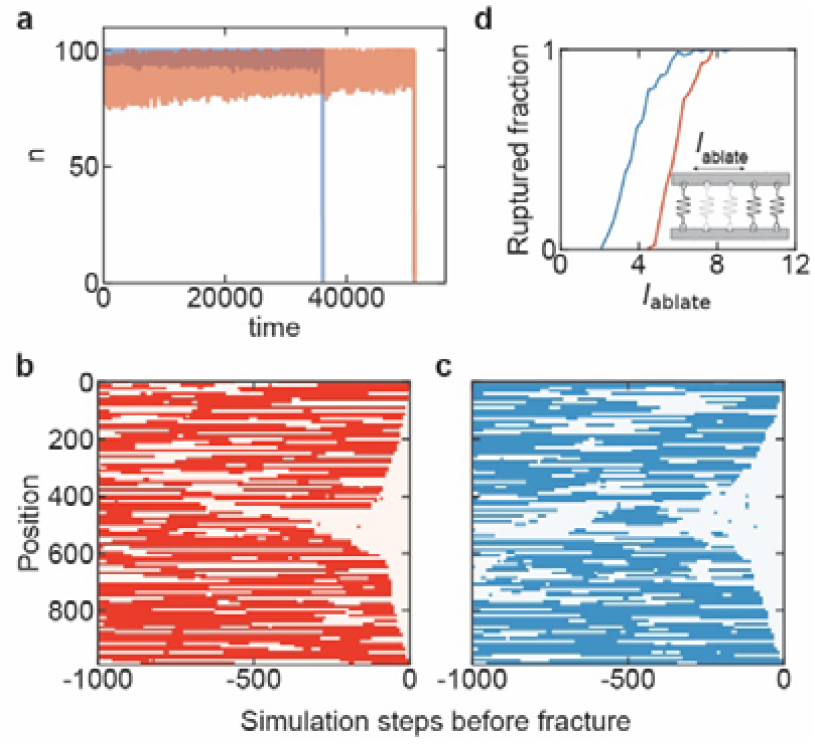
Fracturing in the minimal 1D crosslinker model. **a**, Time trace of the bound number of catch bonds (red) and slip bonds (blue) in a network undergoing a linearly increasing stress (see Extended Data Table 1 for parameters). As the catch bonds have faster dynamics than the slip bonds, a larger spread in the bound fraction is observed. After a long time of steady state fluctuations, the networks suddenly fracture as the number of linkers rapidly goes to 0. **b, c**, Kymographs showing at which positions there are bonds (red for catch bonds, blue for slip bonds) or no bonds (white). At steady state, linkers continuously bind and unbind (−1000 to approximately -300 steps). Cracks can spontaneously initiate and propagate through the network (the last ∼300 steps of the simulation) for both catch and slip bonds in a similar manner. **d**, The fraction of 1D-networks that rupture when a gap of varying ablation length *l*_ablate_ is introduced for both catch (red) and slip bonds (blue). Inset: schematic of the ablation simulation.

**Extended Data Fig. 6:**
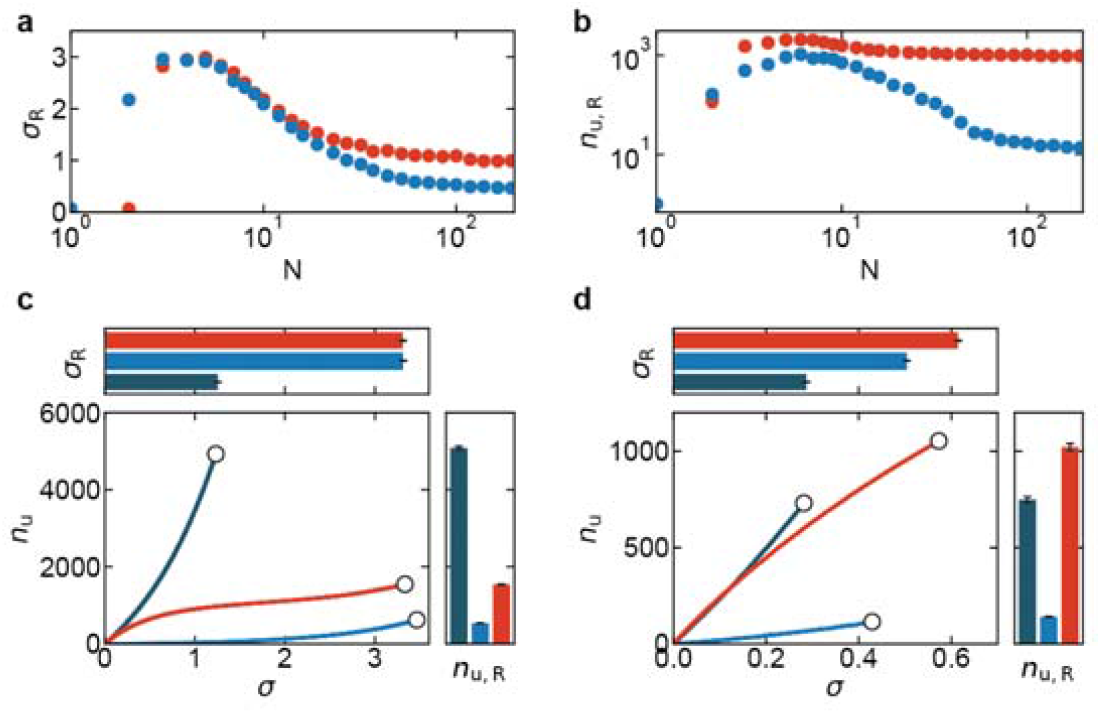
Simulations show that catch bonds only provide a mechanical advantage over slip bonds when they are mobile and present in sufficiently large numbers. The system size dependence of the rupture stress (**a**) and bond turnover at the point of rupture (**b**) reveals that catch bonds (red) are only stronger than slip bonds (blue) for networks larger than ∼10 bonds, emphasizing that the increased network strength by catch bonding is an emergent property (Supplementary Information). Each data point is the average of 10 repeats and the standard errors are smaller than the symbol size. **c**, Catch bond-induced network strengthening is not observed when crosslinkers are immobile and rebind in the same location from which they unbound. The bond turnover as a function of stress reveals catch bonds (red) cause more dynamic materials (right), but do not enhance strength (top) compared to strong slip bonds (light blue) and are less dynamic than networks consisting of weak slip bonds (dark blue). The error bars represent the standard error on basis of 10 repeats per condition. **d**, We also considered a three-state model where linkers are doubly bound, singly bound or unbound (Supplementary Information “Three-state model”). Similar to the two-state model, the bond turnover as a function of stress reveals that networks of catch bonds (red) are stronger and more deformable than networks of strong slip bonds (light blue) or weak slip bonds (dark blue).

**Extended Data Fig. 7:**
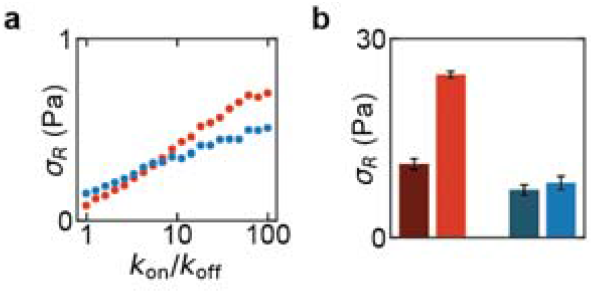
Catch bonding is only effective when the bond lifetime is high. **a**, Simulations of the rupture stress as a function of the bond lifetime 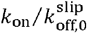, keeping 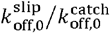 fixed (see Methods and Extended Data Table 1), shows that catch bonds (red) are only stronger than slip bonds (blue) when the binding rate is high. **b**, Consistent with the simulations, enhancing the bond lifetime in experiments by decreasing the temperature from 25 °C (light) to 10 °C (dark) increases the rupture stress more steeply for wild type α-actinin-4 (red) than for K255E (blue).

**Extended Data Fig. 8:**
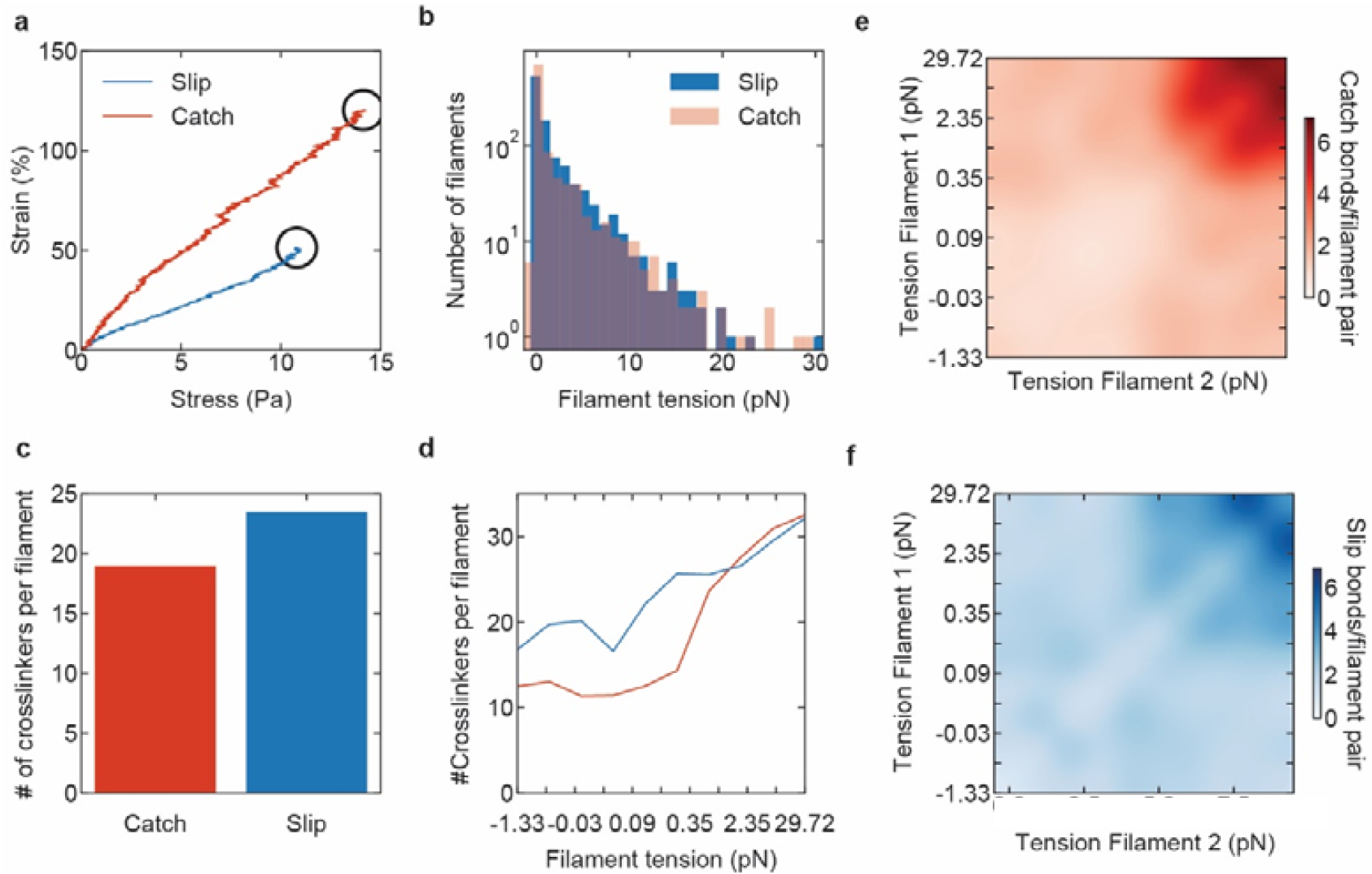
Actin network simulations. **a**, Stress-strain curve of the simulated catch (red) and slip bond (blue) actin networks. The black circles indicate the yielding points of both networks (see methods for details). **b**, Distribution of tension on actin filaments in networks for simulated catch bond (red) and slip bond (blue) with 10 Pa stress. **c**, The polymer network simulation predicts that the average number of active crosslinkers per filament at 10 Pa is lower for catch bonds than for slip bonds, in line with the 1D simulations (panel a) and the catch bond’s lower bond lifetime. **d**, The average number of crosslinkers as a function of the filament tension shows that slip bonds are mainly enriched on low-tension filaments. The tension on the x-axis is binned such that each bin contains 10% of the filaments. **e-f**, Similar plots as Fig. 3d but then for the catch (d, red) and slip bond (e, blue) simulations separately: the active crosslinkers are binned according to tension acting on pairs of filaments (tension on filament 1 on x-axis, tension on filament 2 on y-axis) connected by the crosslinkers. 10×10 bins are used, and the distribution was smoothed using bicubic interpolation. The tension spacing along the x- and y-axis is non-uniform, such that each bin includes 10% of the filaments. These plots show that both catch and slip bonds preferentially connect tense filaments, likely because of geometrical reasons and/or because filament tension resulted from having more crosslinkers bound. However, catch bonds localize more strongly to tense filaments than slip bonds (Fig. 3d) because they bind less to the rest of the network due to their higher off-rate in the absence of force and therefore redistribute to the tense filaments (Fig. 3e).

**Extended Data Fig. 9:**
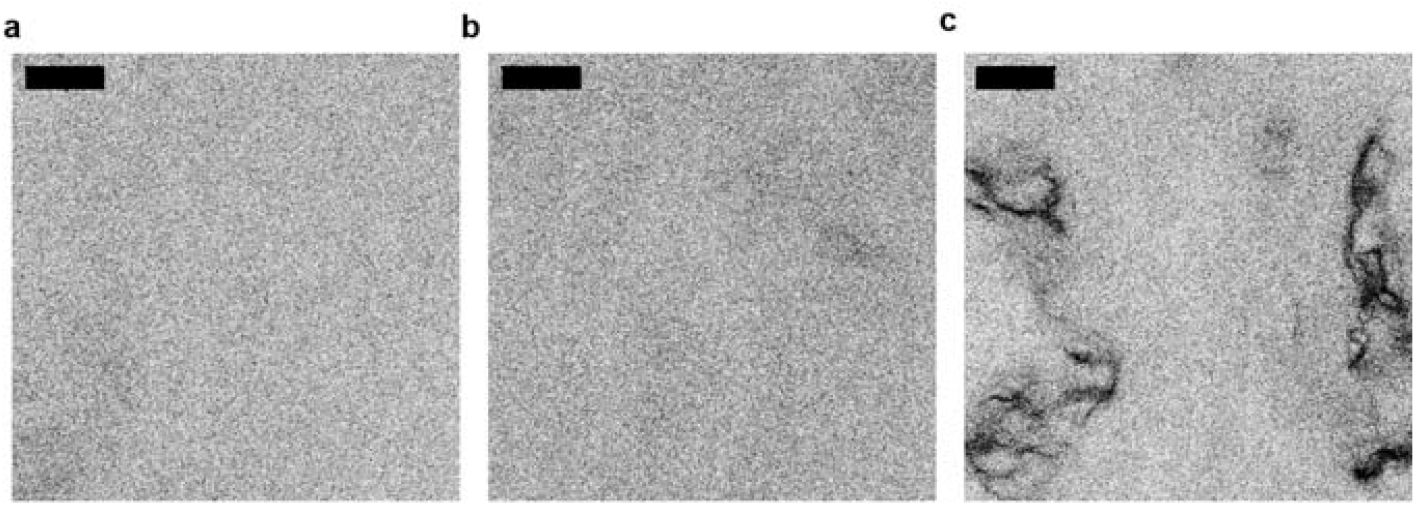
Confocal fluorescence images of crosslinked actin networks. 10% of the actin monomers were labeled with Alexafluor-647. At a 1:100 crosslinker:actin molar ratio, the actin networks studied in this work are isotropic and spatially uniform, for both wild type (**a**) and K255E α-actinin-4 (**b**). We do not observe any discernable structure because the mesh size is ∼200 nm, which is on the order of the diffraction limit, indicating that filaments are isotropically crosslinked rather than bundled. **c**, For comparison, actin bundle clusters were observed at a 1:25 α-actinin-4:actin molar ratio. The color coding was inverted for all images to improve the visual contrast between bundles and background. Scale bars are 20 μm.

**Extended Data Fig. 10:**
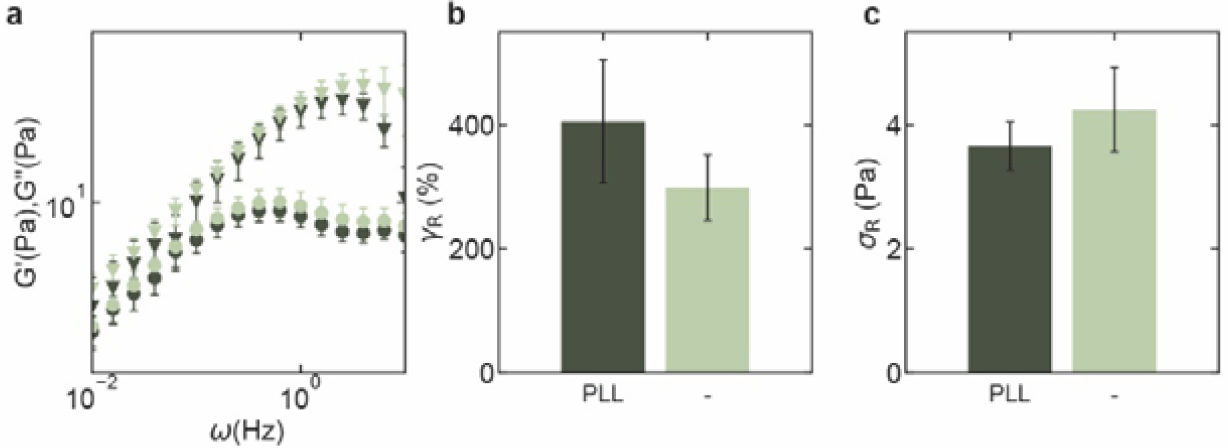
Fracturing occurs within the actin network, not at the rheometer-network interface. The rheology of wild type α-actinin-crosslinked actin networks was compared in the presence (dark green) or absence (light green) of a Polylysine-coated surface on both the bottom and top plate of the rheometer (see Methods section for details). **a)** a frequency sweep at zero prestress shows that the linear rheology is unaffected by changing the rheometer-network interface. The storage (triangles) and loss moduli (circles) were measured as a function of frequency using small amplitude oscillatory shear. b) the network rupture strain and c) rupture stress (bottom) are not significantly affected by the addition of Polylysine at the rheometer-network interface.

**Extended Data Table 1:** Table of parameters used for the minimal 1D crosslinker model. In all simulations, we use 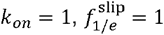 and 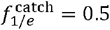. The applied stress (σ), stress rate 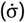, number of crosslinkers (*N*) and bond affinities (*k*_2_ and *k*_2_) varied for different simulations as shown in this table. For catch bonds (red in all graphs) and strong (low temperature) slip bonds (dark blue in all graphs), we used 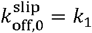. For weak (high temperature) slip bonds (light blue in all graphs), we use 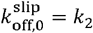. Lastly, 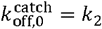 for catch bonds and 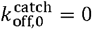 for both strong and weak slip bonds. NA stands for ‘Not Applicable’. All units are dimensionless as explained in the methods.

| Figure(s) | $\sigma$ | $\dot{\sigma}$ | $N$ | $k_1$ | $k_2$ |
| --- | --- | --- | --- | --- | --- |
| Fig. 2d, Extended Data Fig. 6c-d, 8 | NA | $10^{-5}$ | 100 | $10^{-2}$ | $10^{-1}$ |
| Fig. 3a-b | 0.3 | NA | $10^3$ | $10^{-2}$ | $10^{-1}$ |
| Extended Data Fig. 6a-b | NA | $10^{-5}$ | 1 – 100 | $10^{-2}$ | $10^{-1}$ |
| Extended Data Fig. 7a | NA | $10^{-5}$ | 100 | $10^{-2} - 10^1$ | $10 * k_1$ |

**Extended Data Table 2:**
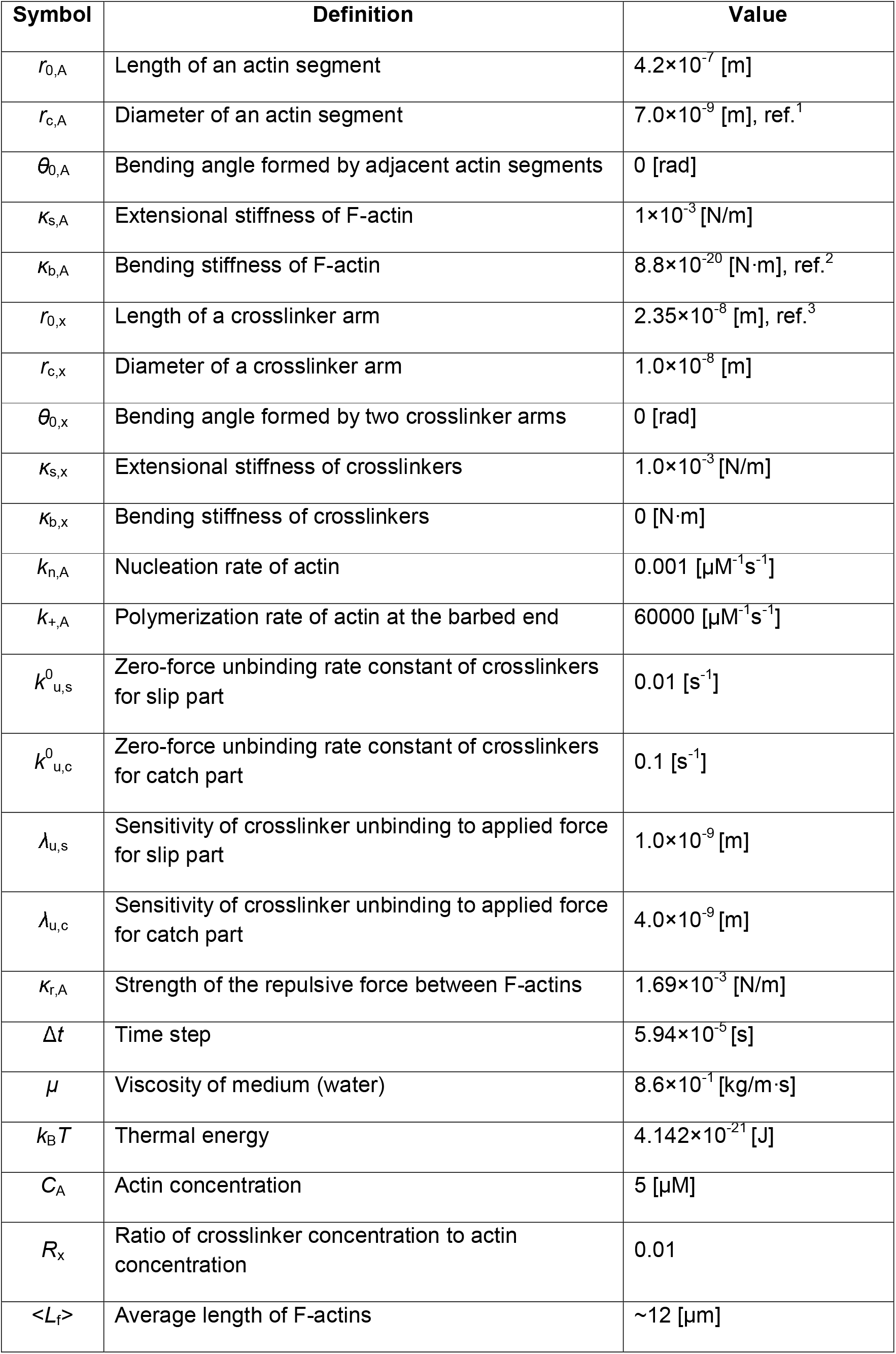
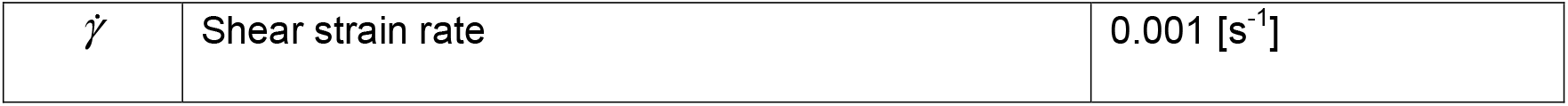
Table of parameters used for the actin network simulations. References are provided for some of the parameters.

| Symbol | Definition | Value |
| --- | --- | --- |
| $r_{0,A}$ | Length of an actin segment | $4.2 \times 10^{-7}$ [m] |
| $r_{c,A}$ | Diameter of an actin segment | $7.0 \times 10^{-9}$ [m], ref.<sup>1</sup> |
| $\theta_{0,A}$ | Bending angle formed by adjacent actin segments | 0 [rad] |
| $\kappa_{s,A}$ | Extensional stiffness of F-actin | $1 \times 10^{-3}$ [N/m] |
| $\kappa_{b,A}$ | Bending stiffness of F-actin | $8.8 \times 10^{-20}$ [N·m], ref.<sup>2</sup> |
| $r_{0,x}$ | Length of a crosslinker arm | $2.35 \times 10^{-8}$ [m], ref.<sup>3</sup> |
| $r_{c,x}$ | Diameter of a crosslinker arm | $1.0 \times 10^{-8}$ [m] |
| $\theta_{0,x}$ | Bending angle formed by two crosslinker arms | 0 [rad] |
| $\kappa_{s,x}$ | Extensional stiffness of crosslinkers | $1.0 \times 10^{-3}$ [N/m] |
| $\kappa_{b,x}$ | Bending stiffness of crosslinkers | 0 [N·m] |
| $k_{n,A}$ | Nucleation rate of actin | $0.001$ [$\mu\text{M}^{-1}\text{s}^{-1}$] |
| $k_{+,A}$ | Polymerization rate of actin at the barbed end | $60000$ [$\mu\text{M}^{-1}\text{s}^{-1}$] |
| $k_{u,s}^0$ | Zero-force unbinding rate constant of crosslinkers for slip part | $0.01$ [$\text{s}^{-1}$] |
| $k_{u,c}^0$ | Zero-force unbinding rate constant of crosslinkers for catch part | $0.1$ [$\text{s}^{-1}$] |
| $\lambda_{u,s}$ | Sensitivity of crosslinker unbinding to applied force for slip part | $1.0 \times 10^{-9}$ [m] |
| $\lambda_{u,c}$ | Sensitivity of crosslinker unbinding to applied force for catch part | $4.0 \times 10^{-9}$ [m] |
| $\kappa_{r,A}$ | Strength of the repulsive force between F-actins | $1.69 \times 10^{-3}$ [N/m] |
| $\Delta t$ | Time step | $5.94 \times 10^{-5}$ [s] |
| $\mu$ | Viscosity of medium (water) | $8.6 \times 10^{-1}$ [kg/m·s] |
| $k_B T$ | Thermal energy | $4.142 \times 10^{-21}$ [J] |
| $C_A$ | Actin concentration | 5 [$\mu\text{M}$] |
| $R_x$ | Ratio of crosslinker concentration to actin concentration | 0.01 |
| $\langle L_f \rangle$ | Average length of F-actins | ~12 [$\mu\text{m}$] |
| $\dot{\gamma}$ | Shear strain rate | 0.001 [s<sup>-1</sup>] |

